# How do Vampires Suck Blood?

**DOI:** 10.1101/2022.10.31.514445

**Authors:** Meng Gou, Xuyuan Duan, Jun Li, Yaocen Wang, Qingwei Li, Yue Pang, Yonghui Dong

**Affiliations:** College of Life Science, Liaoning Normal University, Dalian 116081, China; Lamprey Research Center, Liaoning Normal University, Dalian 116081, China; Department of Life Sciences Core Facilities, Weizmann Institute of Science, Rehovot 7610001, Israel

**Keywords:** Lamprey, Spatial Metabolomics, Blood-sucking, Database

## Abstract

Lampreys are blood-sucking vampires in the marine. From a survival perspective, it is expected that lamprey buccal gland exhibits a repository of pharmacologically active components to modulate the host’s homeostasis, inflammatory and immune responses. Several proteins have been found to function as anticoagulants, ion channel blockers, and immune suppressors in lampreys, while small metabolites have never been explored in detail. In this study, by analyzing the metabolic profiles of 14 different lamprey tissues, we have identified two groups of blood-sucking-associated metabolites, i.e., kynurenine pathway metabolites and prostaglandins, in the buccal gland and they can be injected into the host fish to ensure a steady and sustained blood flow to the feeding site. These findings demonstrate the complex nature of lamprey buccal gland and highlight the diversity in the mechanisms utilized for blood-sucking in lampreys. In addition, a lamprey spatial metabolomics database (https://www.lampreydb.com) was constructed to assist studies using lampreys as model animal. The database contains detailed qualitative, quantitative, and spatial distribution information of each detected metabolite, and users can easily query and check their metabolites of interest, and/or identify unknown peaks using the database.

**Significance Statement:** Lampreys are one of the two surviving jawless vertebrate groups that hold the key to our understanding of the early vertebrate evolution, adaptive immune origin, and developmental neurobiology. Here, we applied a spatial metabolomics approach to study the lamprey-host interaction. Two groups of metabolites, i.e., kynurenine pathway metabolites and prostaglandins, were found in the lamprey buccal gland, which modulate the host’s homeostasis, inflammatory and immune responses. The establishment of the first tissue-wide spatial lamprey metabolomics database in this study facilitate future studies in biochemistry, clinical chemistry, natural product discovery, medicine, and metabolomics using lampreys as a model animal.

## Introduction

Lampreys are primitive vertebrates that, together with hagfishes, represent the only two extant genera of jawless fish known as the Agnatha ^1, 2^. Accumulating fossil evidence has demonstrated that lampreys in the Devonian period are already almost identical to the modern adult lampreys, with well-developed oral disc, annular cartilages, and circumoral teeth ^3–5^, suggesting the evolutionary long-term stability of lampreys.

Lampreys are aquatic, eel-shaped animals. Some species live in freshwater for their entire lives such as the northeast lamprey (*Lampetra morii*), while others, including the sea lamprey (*Petromyzon marinus*) and the Japanese lamprey (*Lampetra japonica*), usually migrate to the sea to feed ^6^. The life cycle of lampreys typically starts in freshwater, where fertilized eggs hatch into small, wormlike larvae (ammocoetes). Ammocoetes differ significantly from their adults in morphology, and they usually burrow into soft substrates where they filter feed on organic matter. Lampreys remain as larvae for about 2-8 years, which appears to be influenced by factors such as climate and food quality ^5, 7^. After metamorphosis, the parasitic phase lampreys migrate into the ocean and feed on the blood and body fluids of the hosts. In the last stage, the non-parasitic phase adult lampreys return to freshwater to spawn and die ^6, 8^.

Forty lamprey species are currently recognized for the extant lampreys, of which 18 species are parasitic ^9^. Almost all blood-sucking animals are invertebrates, such as fleas, ticks, leeches, and mosquitoes, and only lampreys and vampire bats are true ectoparasites ^10^. Because of the blood-sucking habit, lampreys are often referred to as “vampire fish”. Parasitic lampreys usually attach themselves to the body surface of the host through their sucker-like oral disc, rasp a hole in the skin with a tongue-like piston tipped with denticles that form the cutting edges, and suck the blood of the host for days. As such, parasitic lampreys must suppress the immune response (that can lead to itching or pain and thus trigger defensive behavior on their hosts), nociceptive response (that can initiate host defense behavior), and hemostasis (the vertebrate mechanisms that prevent blood loss) of the host to ensure successful and long-term blood feeding. Extensive studies have revealed that the lamprey buccal gland secretes various proteins that function as anticoagulants, ion channel blockers, and immune suppressors ^6, 10, 11^. While small molecules in the secretion have never been explored in detail. Considering their unique phylogenetic position and being one of the only two true ectoparasites, it is expected that lampreys have independently evolved unique metabolites for blood-feeding and parasitism. Detecting and identifying these metabolites will therefore undoubtedly improve our understanding in how lampreys suck blood at metabolic level. In addition, these metabolites may provide new insights into the development of effective drugs in anti-inflammation and pain-relief. To this end, we have performed a spatial metabolomics analysis of 14 different lamprey tissues. The lamprey buccal gland was particularly investigated due to the reason that it is a blood-sucking organ, and that an unexpected rich and unique metabolic profile was detected in buccal gland. Finally, we have constructed a lamprey spatial metabolomics database to facilitate studies in biochemistry, clinical chemistry, natural product discovery, medicine, and metabolomics using lampreys as a model animal.

## Results

### Tissue-wide spatial metabolomics of lamprey

Fourteen lamprey tissues were dissected and subjected to untargeted metabolomics using liquid chromatograph mass spectrometry (LCMS) (Fig. 1a). The raw data quality and inter- batch variation were assessed on 5 pooled quality control (QC) samples obtained at both positive and negative ion modes using R package RawHummus ^12^. This software randomly selects 6 ion peaks evenly across the entire retention time range and adopts 12 metrics that are closely related to LC peaks shape, MS1 (full scan spectra), and MS2 (fragment ion spectra) for rapid and comprehensive data quality evaluation. The resulting reports showed that the maximum mass accuracy variation was less than 2 ppm, the maximum retention time (RT) shift was less than 0.2 min, and the ion intensity coefficient of variation (CV) was less than 10% at both positive and negative ion modes, suggesting an excellent reproducibility in our study (File S1 and S2). In addition, an internal standard (IS), 2-chloro-L-phenylalanine, was used for rapid inter-batch variation evaluation, and the result confirmed a good reproducibility within the samples (peak area variation of the IS was less than 10% across all the samples at both positive and negative ion mode) (Fig. 1c).

**Fig. 1.**
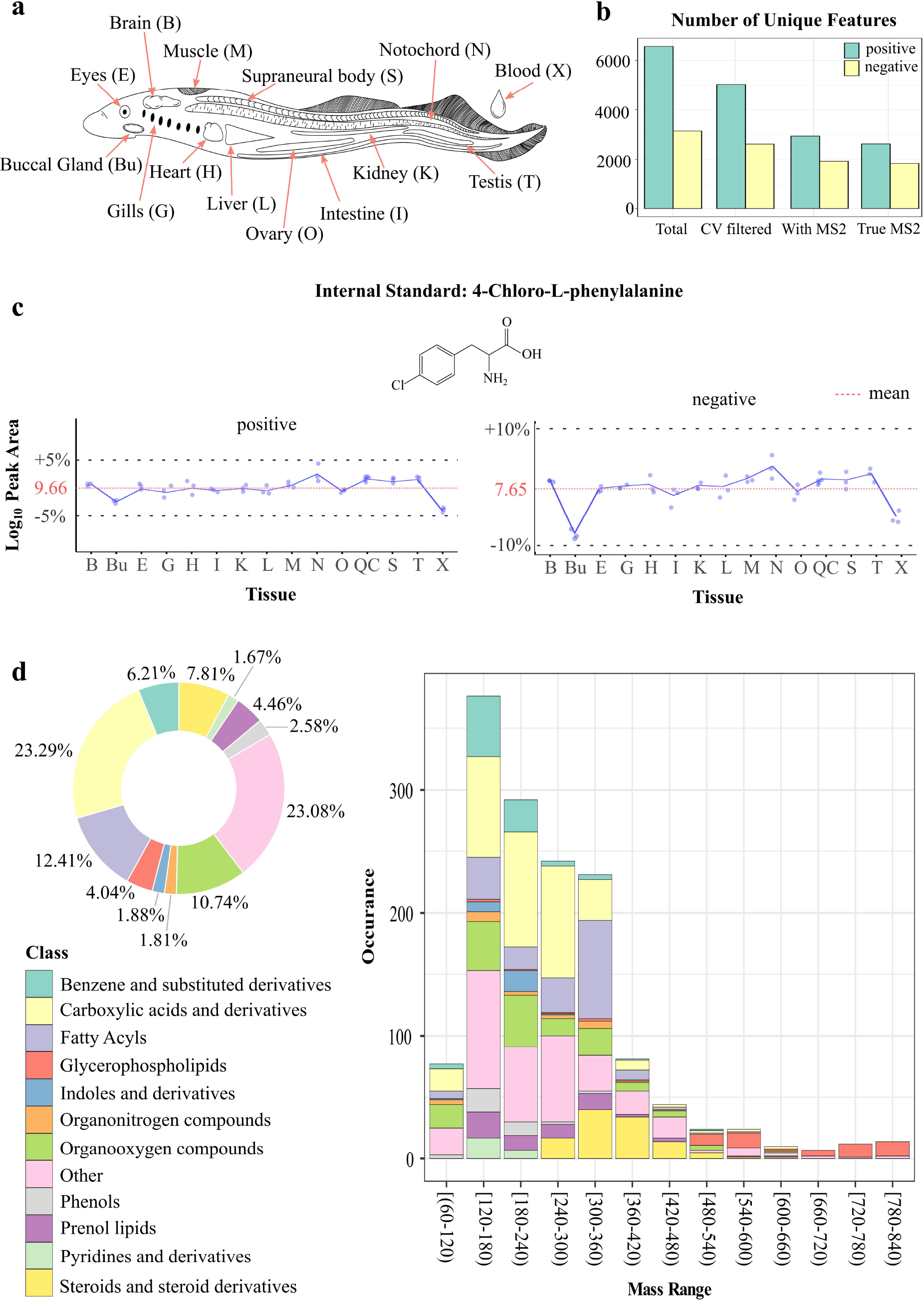
Overview of spatial metabolomics of lampreys. **a**, Anatomical illustration of the 14 lamprey tissues subjected to metabolomics analysis. Each tissue has a unique abbreviation and is kept consistent throughout the figures. **b**, Bar plot showing the number of unique mass features before and after data cleaning at positive and negative ion modes. **c**, Line plots showing the log10 transformed peak area variations of internal standard, 4-chloro-L- phenylalanine, detected at positive and negative ion modes, respectively (n = 3 for each tissue). **d**, Summary of the total identified metabolites in different lamprey tissues. Pie chart showing the percentage of identified metabolites belonging to different chemical classes. Histogram showing the distribution of chemical classes along the mass range. Each chemical class is represented by a unique color code.

Raw data were preprocessed using Compound Discoverer (v.3.2, Thermo Scientific) software, and ion peaks were grouped based on both RT, adducts, and isotope deconvolutions to generate “unique mass features” (i.e., ion species derived from the same metabolite such as adducts, isotopes, and in-source fragments were grouped as one). In total, 6,568 and 3,143 unique features were detected in positive and negative ion modes, respectively (Fig. 1b). A final data cleaning was performed by removing mass features with CV > 30% in the QC sample group, features without MS2 spectrum (i.e., no MS/MS fragmentation was performed on that precursor ion), and features with wrong MS2 spectrum (i.e., wrong precursor ion was assigned for MS/MS fragmentation). The resulting 2621 (positive ion mode) and 1835 (negative ion mode) mass features were left for subsequent metabolite identification and statistical analysis (Fig. 1b).

Tentative and putative metabolite annotations were performed with Compound Discoverer based on accurate mass measurements (<5 ppm error), isotope distribution similarity, and manual assessment of fragmentation spectrum matching (when applicable) against our in-house database (∼3600 standards) and mzCloud database (https://www.mzcloud.org). Additional metabolite identifications were made by searching against the Human Metabolome Database (https://hmdb.ca) ^13^ and Lipid Map (https://www.lipidmaps.org) ^14^ using Progenesis QI (v.2.3, Waters), and all public MS/MS positive and negative databases using MS-DIAL (v4.0) ^15, 16^. This step led to the detection of a large diverse of metabolite classes in lamprey; among them, carboxylic acid and its derivatives are the most abundant class (Fig. 1d).

### The blood-sucking organ buccal gland is a metabolic outlier tissue

Principal component analysis (PCA) was applied for initial examination of the metabolic profiles of the different lamprey tissues. Surprisingly, the PCA score plots from both positive and negative ion mode data revealed that the buccal gland was far separated from all the other tissues (Fig. 2a), suggesting that buccal gland has a distinct metabolic profile. Hierarchical clustering heatmap showed that one-third of the mass features were highly abundant in buccal gland in positive ion mode (Fig. 2b), and more than half of the features were enriched in buccal gland in negative ion mode (Fig. 2b). Further statistical analysis showed a significant accumulation of metabolites in buccal gland. For instance, 127 and 182 mass features were found over 1000-times higher in buccal gland compared to all the other 13 tissues (FDR-adjusted p-value < 0.001) in positive and negative ion modes, respectively. It is worth noting that it is typically not straightforward to directly compare the metabolic profiles of different tissues due to the differences in tissue-specific matrix effects ^17, 18^. However, the equal detection of IS in different tissue groups may indicate that the tissue- specific matrix effects differences were not significant in our study (Fig. 1c). Nevertheless, our objective here was not to identify biomarkers to distinguish the buccal gland from any other lamprey tissues. Instead, here we would like to fish out lamprey buccal gland-specific metabolites.

**Fig. 2.**
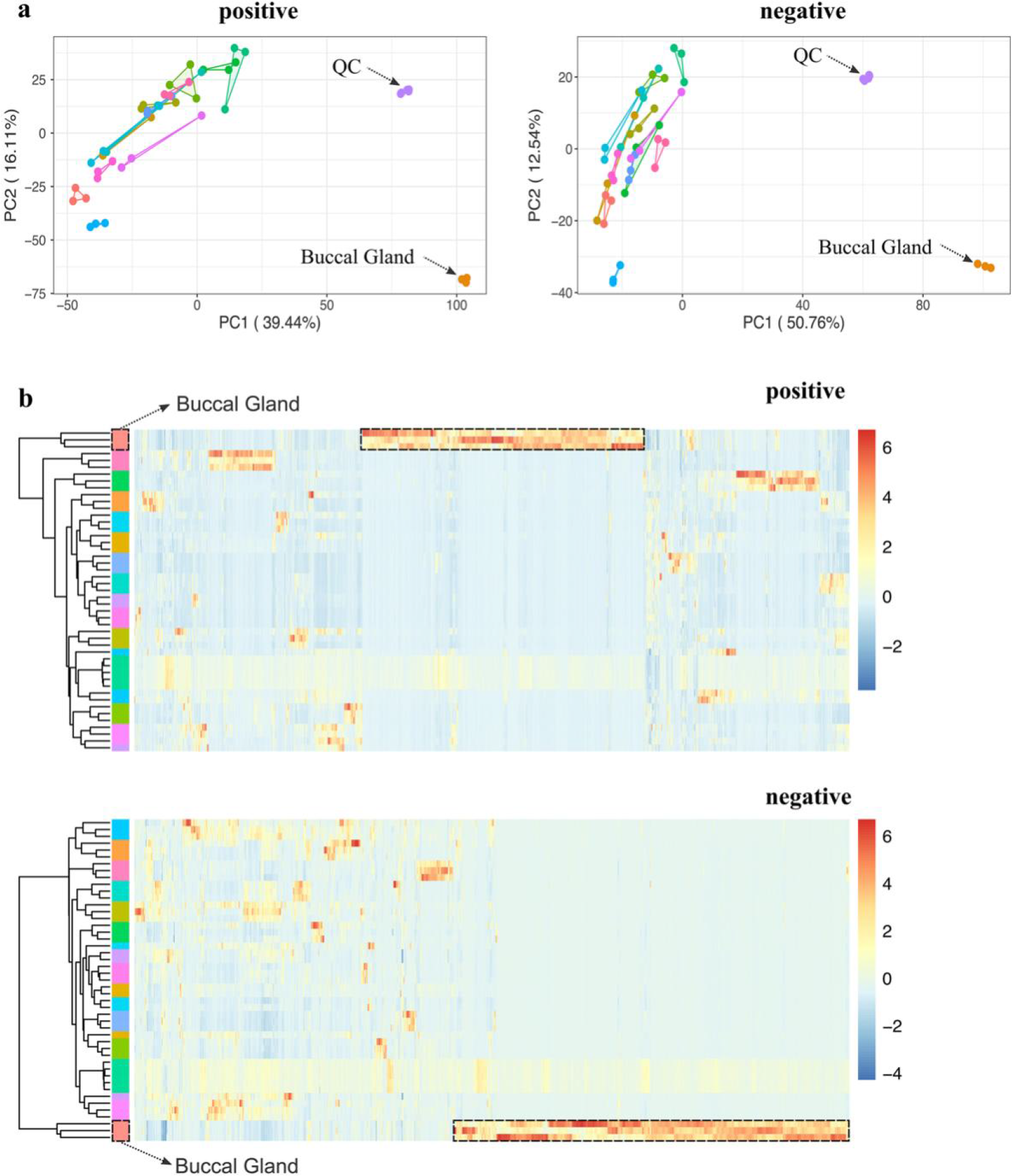
Unsupervised multivariate analysis of the metabolic profiles of 14 different lamprey tissues. **a**, Principal component analysis (PCA) score plots of the metabolic profiles acquired at positive and negative ion modes, respectively (n = 3 for each tissue except that n = 5 for QC samples). **b**. Hierarchical clustering heatmaps of the metabolic profiles acquired at positive and negative ion modes, respectively (n = 3 for each tissue except that n = 5 for QC samples). Each row represents a sample, and each column represents a metabolite. Each colored cell (red higher, blue lower) on the map corresponds to a z-score normalized mass feature.

Parasitic lampreys are known to secrete proteins that possess anticoagulant and vasodilator activities from the buccal glands during feeding on their hosts. Insensitive studies have been performed to detect, identify and study the functions of those proteins ^6, 8, 10, 11^. In contrast, there have been surprisingly few studies on small molecules in lamprey buccal glands ^19^. Considering the extremely unexpected rich and unique metabolic profiles of the buccal gland and its important biological function as a blood-sucking organ, we have therefore focused on the buccal gland in the subsequent studies to investigate the lamprey blood-sucking mechanism at the functional metabolic level.

### Tryptophan-kynurenine pathway metabolites and prostaglandins were exclusively accumulated in the buccal gland

In total, more than 1,500 mass features were found highly abundant in lamprey buccal gland (FC >= 10 and FDR-adjusted p-value < 0.05). Among them, 272 were tentatively identified and they belong to over 30 different chemical classes, such as fatty acyls, steroids, and steroid derivatives. These buccal gland-specific mass features are perfect candidates for screening blood-sucking associated metabolites. Notably, a complete kynurenine pathway (KP) was detected in the buccal gland (Fig. 3a). The MS/MS spectrum of each KP pathway metabolite, annotation of their major fragments, and head-to-tail library match plots were shown in Figure S1-6. As clearly shown in the anatomical heatmap, most of the KP metabolites were exclusively accumulated in buccal gland (Fig. 3a-b). For instance, N- formylkynurenine was found between 229.0-14676.9 times higher in buccal gland compared to all the other 13 tissues, and kynurenine was between 27355-46627.6 times higher in buccal gland (Fig. 3a). In addition, a lamprey buccal gland-specific KP pathway metabolite, namely 3-hydroxykynurenine-O-sulfate ^19^, was also identified with its fold change values ranging from 2713.2 to 47791.6 in buccal gland compared to other tissues (Fig. 3a). Although its function is still unclear, the detection of 3-hydroxykynurenine-O-sulfate in other blood- sucking insects, such as *Rhodnius prolixus* ^20^, suggest that it might be a blood-feeding related metabolite. The KP is rate-limited by its first enzymes, tryptophan 2,3-dioxygenase (TDO) and indoleamine 2,3-dioxygenase (IDO), which convert tryptophan into N-formylkynurenine ^21, 22^ (Fig. 3a). The expression levels of the two major genes were studied by real-time quantitative PCR (qPCR), and the result showed that TDO was highly expressed in the buccal gland while IDO was mostly in the liver (Fig. 3c).

**Fig. 3.**
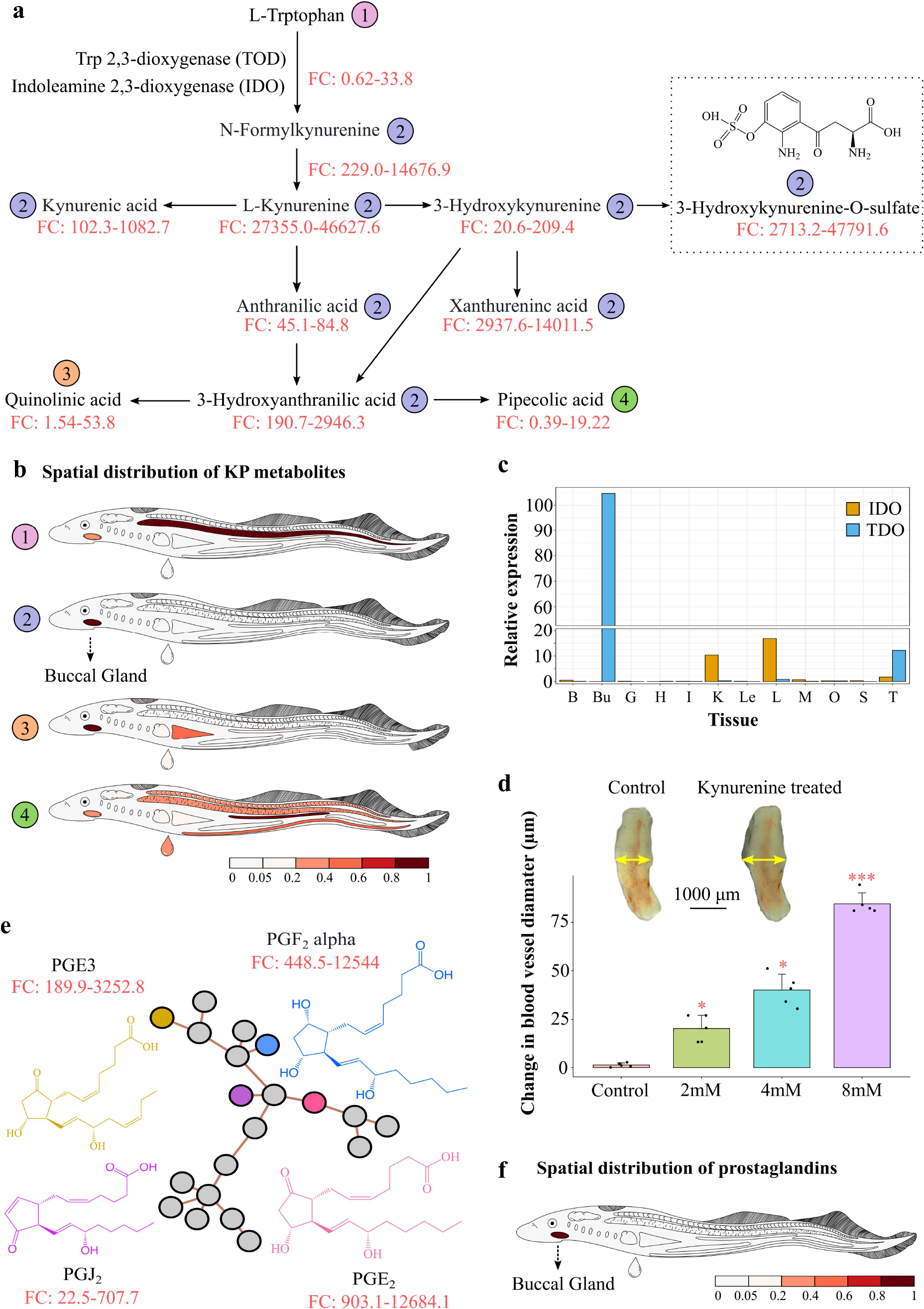
Kynurenine pathway metabolites and prostaglandins detected in lamprey buccal gland. **a**, Schematic representation of the kynurenine pathway (KP). The fold change (FC) value was calculated for each metabolite in the pathway by dividing the peak area of that metabolite detected in the buccal gland by that in all the other lamprey tissues (QC samples were excluded). The FC value range was then displayed underneath each metabolite name. The chemical structure for 3-hydroxykynurenine-O-sulfate, a lamprey-unique KP metabolite was shown in the pathway. **b**, Four unique spatial distribution patterns were detected for KP metabolites. L-tryptophan was accumulated in the notochord (circled in purple); quinolinic acid was mostly found in lamprey buccal gland and liver (circled in orange); pipecolic acid was more abundant in kidney (circled in green); and all the rest KP metabolites were exclusively accumulated in lamprey buccal gland (circled in blue). **c**, the relative expression of tryptophan 2,3-dioxygenase (TOD) and indoleamine 2,3-dioxygenase in different lamprey tissues. The tissue abbreviations are the same as shown in Fig. 1a. **d**, Bar plot showing the changes in catfish blood vessel diameter upon treatments of different concentrations of kynurenine. The insert representative figures are the optical images of catfish blood vessels before and after kynurenine treatment. Asterisks denote significant differences (*, p < 0.05; ***, p < 0.001; Wilcoxon signed-rank test) between kynurenine**-**treated and PBS treated (control) blood vessels. **e**, A sub molecular network of prostaglandins was detected in lamprey buccal gland. Chemical structures of the 4 identified prostaglandins were displayed. The fold change (FC) value range of each prostaglandin was calculated in the same way as described in Fig. 3a. Grey node represents unidentified mass peaks or in-source fragments of the identified metabolites. **f**, A representative anatomical heatmap showed that all the 4 prostaglandins were exclusively localized in lamprey buccal gland.

The KP has received increasing attention nowadays as its connection to inflammation, the immune system, and neurological conditions is becoming more apparent. These unique roles of KP are potentially associated with lamprey blood-sucking. In addition, the report of 3-hydroxykynurenine-O-sulfate in other blood-sucking species makes KP even more attractive in investigating their roles in lamprey blood feeding. L-Kynurenine, in particular, has been reported to relax the blood vessels, decrease vascular resistance and improve blood flow by lowing the blood pressure ^23, 24^. We, therefore, asked if L-kynurenine could also relax the blood vessels of the host fish. As such, we performed a vasomotor reactivity test on the catfish (*Silurus asotus*) aortic ring. As shown in Fig. 3d, compared to the control group (PBS- treated), the blood vessel diameter significantly increased 15 min after the addition of L- kynurenine, and the degree of increase appeared to be associated with the concentrations of L-kynurenine (Fig. 3d).

Apart from KP metabolites, four prostaglandins (PG), namely PG<u>J</u>2, PGF2alpha, PGE2, and PGE3, were identified in the buccal gland using molecular networking analysis (Fig. 3e). The MS/MS spectrum of each prostaglandin, annotation of their major fragments, and head- to-tail library match plots were shown in Figure S7-10. Anatomical heatmap showed that all the 4 prostaglandins were exclusively accumulated in lamprey buccal gland (Fig. 3f). For instance, PGE2 was found between 903.1-12684.1 times higher in the buccal gland compared to all the other tissues (Fig. 3e). PGs are a group of well-studied blood-sucking related metabolites with vasodilatory and immunomodulatory capabilities, and they have been discovered in the saliva of many bloodsuckers such as tick ^25–28^, salmon louse ^29, 30^ and forest leech ^31^. It is well established that there is great similarity in the salivary components among different bloodsuckers, and one consistency is the presence of PGs. As such, PGs were selected as another potential blood-sucking related metabolite in lamprey for further studies.

### Tryptophan-kynurenine pathway metabolites and prostaglandins can be transferred from lampreys to their host fish

During blood-sucking, parasitic lampreys attach to the host fish by their oral disks, penetrate the skin through the action of their toothed tongue-like pistons, create a feeding niche at the bite site, and then start feeding ^32^. On the other side, host defenses will attempt to counterattack and stop lamprey feeding at the bite site. To ensure continuous blooding feeding, lampreys secrete a plethora of components and inject them into the hosts to suppress host hemostasis, inflammation, and immunity.

In this study, we have identified two groups of metabolites, i.e., KP metabolites and prostaglandins, in the lamprey buccal gland that may play significant roles in lamprey blood- feeding. To verify that these metabolites indeed participated in lamprey blood feeding, we must make sure that they can be secreted from the lamprey buccal gland and injected into the feeding site of host fish. As such, we have designed another set of experiments and performed a “targeted analysis” to monitor changes in the two groups of metabolites during lamprey blooding feeding. It is important to note that the non-parasitic adult Japanese lampreys (*Lampetra japonica*) used in our study already stop feeding, thus the northeast lampreys (*Lampetra morii*) which is a life-long parasite was used to attack host fishes and suck blood ^8^. In brief, sixty lampreys were randomly divided into six groups (10 in each group). The buccal glands of three lamprey groups were dissected, and their secretion was collected before blood-sucking. Another three groups of lampreys were fed with catfish (*Silurus asotus*) for 20 min, and then their buccal gland secretion was collected. In addition, three sampling sites from the catfish, i.e., the blood-sucking site and two non-blood-sucking sites of the host, were collected. In total, the LCMS analyses include 5 sample groups: group 1 is the buccal gland from lamprey before blood-sucking; group 2 is the blood-sucking site from catfish; groups 3 and 4 are none-blood-sucking sites from catfish, and group 5 is the buccal gland from lamprey after blood-sucking (Fig. 4a). By comparing between group 1 and group 5, we are able to know that if the amounts of these metabolites are reduced in buccal gland after blood-sucking; and by comparing among group 2, 3, and 4, we could check if these metabolites are transferred from lamprey buccal gland to the sucking site of the host fish.

**Fig. 4.**
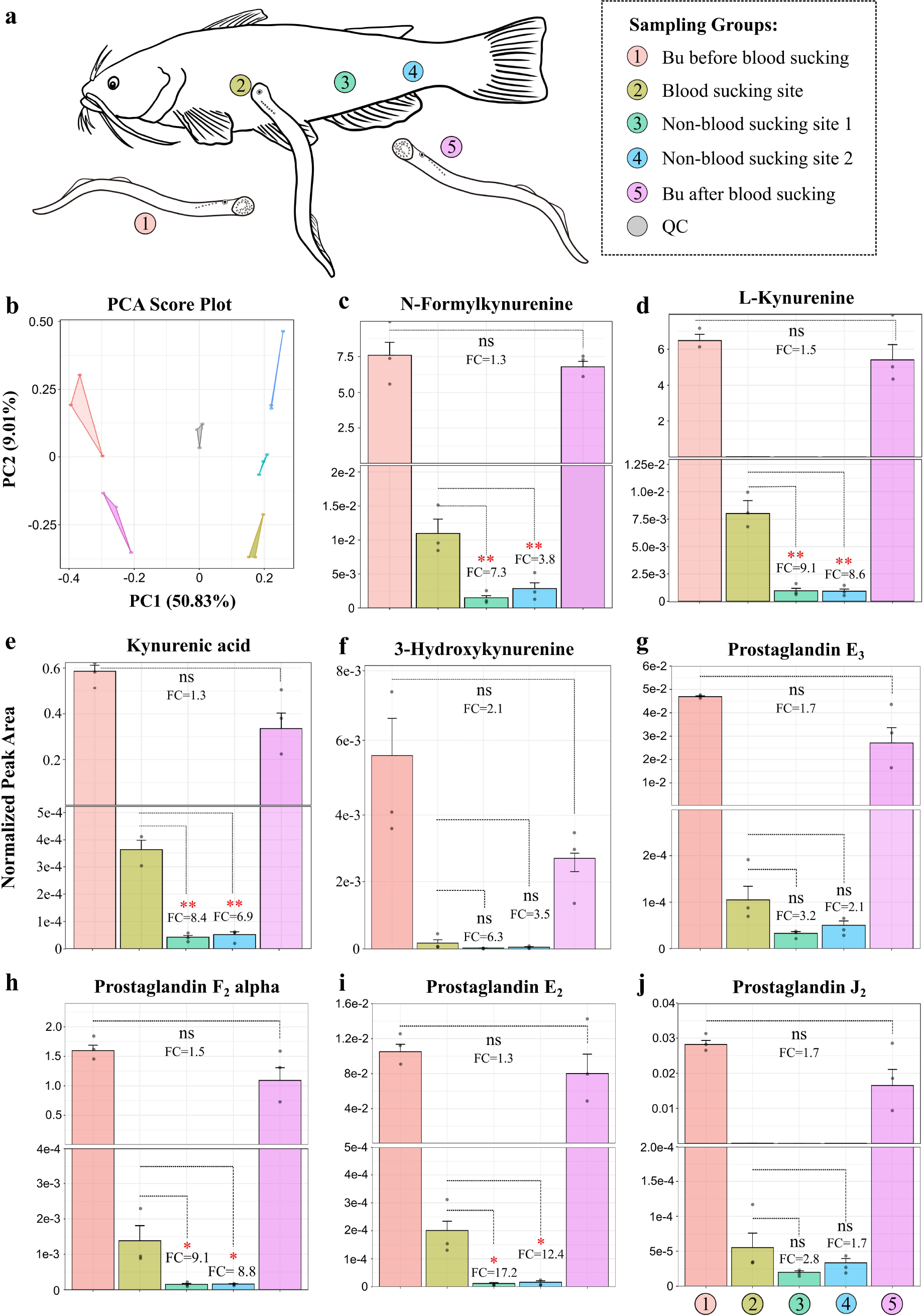
Experimental design for lamprey blooding sucking. **a**, Five groups of lampreys were used for blood-sucking experiment. Group 1 is the buccal gland collected from lampreys before blood-sucking; Group 2 is the feeding-site from the host fish (catfish); Groups 3 and 4 are non-feeding sites from the host fish; and group 5 is the buccal glands collected from lampreys after blood-sucking. **b**, Principal component analysis (PCA) score plots of the metabolic profiles of the 5 sample groups (n = 3 for each group). **c-j**, Measurement of relative peak areas of N-formylkynurenine (c), L-Kynurenine (d), kynurenic acid (e), 3-hydroxykynurenine (f), prostaglandin E3 (g), prostaglandin F2 alpha (h), prostaglandin E2 (i) and prostaglandin J2 (j). The data are shown as the mean ± SD (n = 3). Asterisks denote significant differences (ns, not significant; *, p < 0.05; **, p < 0.01).

PCA score plot of the LCMS data showed that the 5 sample groups can be well separated, suggesting they have very different metabolic profiles (Fig. 4b). The results for KP metabolites, i.e., N-formylkynurenine, L-kynurenine, kynurenic acid, and 3- hydroxykynurenine, are displayed in Fig. 4c-f. Although no significant statistical differences of the four KP metabolites were observed between group 1 and group 5 (FDR-adjusted p- value > 0.05), fold change analysis showed that the amounts of all the four metabolites were reduced after blood sucking (Fig. 4c-f), suggesting that these metabolites were released from lamprey buccal gland during blood sucking. By contrast, significant statistical differences of three KP metabolites, i.e., N-formylkynurenine, L-kynurenine, and kynurenic acid, were found between group 2 and group 3, and between group 2 and group 4 (FDR-adjusted p- value < 0.05) (Fig. 4c-e). Fold change analysis showed that all the four metabolites were highly accumulated in the blooding sucking site (group 2) compared to non-blood-sucking sites of the host fish (groups 3 and 4), demonstrating that the four KP metabolites were transferred from lamprey buccal gland to the sucking site of the host fish. Similarly, the results for another four KP metabolites, i.e., 3-hydroxykynurenine-O-sulfate, anthranilic acid, xanthurenic acid, and 3-hydroxyanthranilic acid, also confirmed that they can be secreted from the buccal gland and injected into the sucking site of catfish (Fig. S11). Same results were also observed for PGs that albeit no significant statistical differences were found between buccal glands before and after blood-sucking (FDR-adjusted p-value > 0.05), the amount of the four PGs were all reduced in buccal gland after blood-sucking (Fig. 4g-j). The results also showed that all the four PGs increased in the sucking site of catfish compared to those in the none-blood-sucking sites. In particular, PGF2 alpha and PGE2 were statistically higher in the blood-sucking site compared to the non-blood sucking sites (Fig. 4h-i).

### Lamprey spatial metabolomics database

Due to its unique status in vertebrate evolution, lampreys have become an important animal model in diverse research fields ^2, 33^, such as vertebrate evolutionary and development ^7, 34, 35^, fundamental aspects of vertebrate neurobiology ^34, 36^, adaptive immunity ^37, 38^, blood clogging ^39^, and bioactive compound identification ^19^. Metabolomics, as a relatively new member in the “omics” field, provides another powerful tool for lamprey studies. However, the lamprey- specific metabolomics database is still missing. As such, we have established a tissue-wide spatial lamprey metabolomics database, named LampreyDB (https://www.lampreydb.com), using all the identified and annotated metabolites from our experiment. LampreyDB is hosted on Microsoft Azure cloud service, and it is freely accessible. It leverages all the benefits provided by PaaS (platform-as-a-Service) solutions particularly designed to guarantee high levels of security and performance. LampreyDB allows users to explore lamprey-specific metabolites with text-based searches, i.e., chemical formula, *m/z* value, or a list of MS/MS fragments (Fig. 5a). The resulting summary page displays a table containing all matched metabolites from the database. Each metabolite entry is hyperlinked to an individual metabolite description page that contains the following information (if available): metabolite name, class, chemical formula, retention time, accurate *m/z* value, SMILES, inChiKey and chemical structure (Fig. 5b). Interactive MS2 spectrum plot and interactive anatomical heatmap are provided in the same page for each metabolite so that the user can easily explore the fragment peaks of each metabolite, and visually inspect and compare its spatial distribution. Currently, LampreyDB contains information on over 1000 metabolites (2031 records from both positive and negative ion modes) detected in lamprey.

**Fig. 5.**
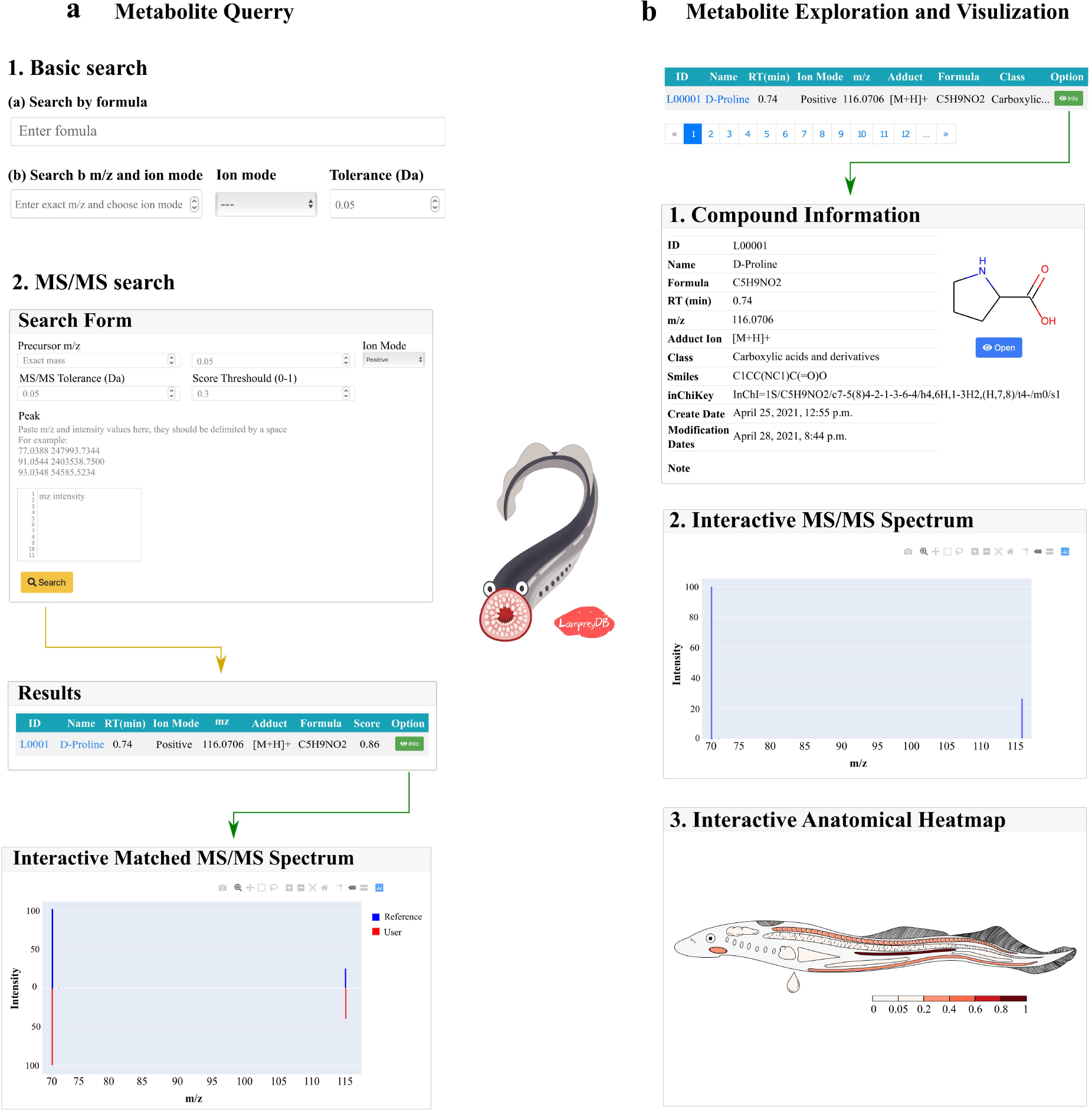
Overview of the LampreyDB spatial metabolomics database. **a**, The LampreyDB allows text-based search such as using chemical formula, *m/z* value, and a list of MS/MS fragments. **b**, The LampreyDB provides rich information including metabolite name, class, chemical formula, retention time, accurate *m/z* value, SMILES, inChiKey, chemical structure, interactive MS/MS spectrum and interactive anatomical heatmap for each metabolite in the database.

## Discussion

When attempting to feed on their hosts, lampreys are challenged by the host defenses, such as hemostasis, inflammation, and immunity. Accordingly, to ensure successful and continuous blood-sucking, lampreys have evolved a complex and sophisticated cocktail of buccal gland secretion components, consisting of a variety of bioactive proteins, peptides, and metabolites, to counteract the host responses. Recent advances in genomics, transcriptomics, and proteomics have allowed the discovery of diverse proteins having anticoagulant and vasodilator activities from the lamprey buccal gland ^6, 8^. While studies on metabolites in the lamprey buccal gland are still missing.

In this study, we applied a spatial metabolomics approach to search for lamprey blood-sucking-related metabolites. Spatial metabolomics is a field of omics research focused on detecting and interpreting metabolites in the spatial context of cells, tissues, organs, and organisms ^40^. Mass spectrometry imaging (MSI) is one of the most noteworthy techniques in spatial metabolomics studies, with the achievable spatial resolution down to cellular and sub- cellular levels to date ^41, 42^. While the molecular coverage is typically low and metabolite identification is challenging in MSI ^43–45^. By contrast, although limited in spatial resolution, the LCMS-based spatial metabolomics approach provides unprecedented sensitivity and molecular coverage, thus allowing a more detailed investigation of the biological system. As our objective is not to map the spatial distribution of metabolite in lampreys, we decided to use LCMS to perform a tissue-wide spatial metabolomics analysis. This approach allows us to explore tissue-specific metabolites in an “untargeted manner”. For instance, by comparing the metabolic profiles among different lamprey tissues, we were able to pinpoint lamprey buccal gland-specific metabolites. These metabolites are perfect candidates for screening lamprey blooding-feeding-related metabolites. With this approach, we were able to detect over 1,500 unique mass features that were highly accumulated in lamprey buccal gland (FC >= 10 and FDR-adjusted p-value < 0.05). Our result implies that the buccal gland contains a much broader complexity of small metabolites than previously anticipated. Further statistical analysis, literature search, and biological function analysis have led to the identification of two groups of candidate metabolites, i.e., the kynurenine pathway (KP) metabolites and prostaglandins (PGs), that may involve in lamprey blood-sucking.

The KP has been a subject of intense research activity in recent years with respect to the discovery of new and important roles its metabolites play, particularly in neuronal and immune function ^21^. In addition to protein synthesis, tryptophan is the precursor of many physiologically important metabolites, whereas over 95 % of tryptophan goes into KP ^21, 46^. To begin with, tryptophan is converted into N-formylkynurenine through the action of either tryptophan 2,3-dioxygenase (TDO) or indoleamine 2,3-dioxygenase (IDO1 and 2), and then into kynurenine by N-formylkynurenine formamidase (FAM). Kynurenine is metabolized mainly by hydroxylation to 3-hydroxykynurenine by kynurenine monooxygenase (KMO) followed by hydrolysis of 3-hydroxykynurenine to 3-hydroxyanthranilic acid by kynureninase (Fig. 3a). KP is highly regulated in the immune system, where it promotes immunosuppression in response to infection or inflammation. Although IDO and TDO share similar functions as a mediator of tryptophan degradation, their biochemical properties are rather different. IDO is a monomeric enzyme less expressed in normal tissues and upregulated during inflammation to suppress immune reaction. In particular, IDO1 has a broad spectrum of activity on immune cells regulation, which controls the balance between stimulation and suppression of the immune system at sites of local inflammation that is relevant to a wide range of autoimmune and inflammatory diseases ^22^. While TDO is a tetramer, and it is highly expressed in liver where it degrades excesses of dietary tryptophan and produces immunosuppressive KP metabolites ^21^. Unlike reported in humans and many other animal species that KP and its first enzyme TDO exist mainly in the liver ^21^, our spatial metabolomics results disclosed that KP and TDO were exclusively located in lamprey buccal gland (Fig. 3b-c). The identification of KP in lamprey buccal gland may imply a unique mechanism for lamprey blood-feeding and parasitism: instead of evolving novel molecules, lampreys accumulate abundant KP metabolites in their buccal gland, and during blood- sucking these metabolites are injected into the sucking site of the host fish, serving as immune suppressors to facilitate continuous blood-sucking. Unlike TDO which mainly existed in lamprey buccal gland, IDO was found mostly expressed in lamprey liver, and its expression level in buccal gland was negligible (Fig. 3c). Nevertheless, the main theory underlying the immunosuppressive function for these enzymes is associated with their canonical tryptophan catabolic properties: TDO and IDO-mediated depletion of tryptophan and the accumulation of kynurenine and other KP metabolites leads to the suppression of immune effector cells and the upregulation, activation, and/or induction of tolerogenic immune cells ^47^. In addition to functioning as immune suppressor, our results demonstrated that kynurenine can also relax the blood vessels of the host fish, thus improving the blood flow of the host fish to towards the blood-sucking site (Fig. 3d). KP is a well-studied pathway, and the structures and functions of its pathway metabolites have been elucidated. In this study, we have also detected a rarely reported KP metabolite, namely 3-hydroxykynurenine O- sulfate, in lamprey buccal gland (Fig. 3a). This metabolite was first identified in lamprey by Odani in 2012 ^19^, but it has been also found in other blood-sucking insects, such as *Rhodnius prolixus* ^20^, suggesting that it is a blood-feeding related metabolite.

Prostaglandins (PGs) are eicosanoid components derived from arachidonic acid through three sequential enzymatic reactions. They act as ‘local hormones’ and regulate a plethora of physiological processes ^31, 48^. PGs are a group of well-studied blood-sucking related metabolites, and they have been discovered in many bloodsuckers, including ticks ^25–28^, salmon louses ^29, 30^ and forest leeches ^31^. Among them, PGE2 is the most commonly found PGs in the secretions, and it has demonstrated prolonged parasitic feeding (anticoagulant), increased blood flow to the site of attachment (vasodilation), and/or evasion of host immune responses (immunomodulator). Another important function of PE2 is to mobilize Ca2^+^ and stimulate the secretion of anticoagulant proteins during the blooding feeding ^49^. It has been also reported that PGE2 could inhibit wound healing by recruiting fibroblasts to the feeding lesion ^27^. In addition to PGE2, PGF2 alpha was also commonly detected in bloodsuckers with reported functions of vasodilation, platelet aggregation inhibition, anti-inflammation, and pain alleviation ^25, 31^. In this study, we have detected another two PGs, i.e., PGJ2 and PGE3, in lamprey buccal gland. Both PGJ2 and PE3 are known to have anti-inflammatory property ^50^. Although they were not reported in other blood-sucking species, it is tempting to speculate a function of the two PGs in blood-feeding and parasitism. Interestingly, all the four PGs were exclusively accumulated in lamprey buccal gland (Fig. 3f), while its precursor arachidonic acid was found in most tissues (Fig. S12). The detection of PGs in lampreys suggested that lampreys may share a similar blood-sucking strategy as other bloodsuckers.

In this study, we were mainly interested in blood-sucking related metabolites in lampreys. While it is known that lampreys are rich sources of a large variety of metabolites with various biologically and physiologically important functions. For instance, we have identified a sulfated bile acid in this study, namely petromyzonol sulfate, known as lamprey- specific sex pheromone ^51, 52^, and its distributions appear to be particularly tissue-specific (Fig.S13). This spatially resolved metabolic information is of great benefit to many studies. As such, we have created a lamprey spatial metabolomics database (https://www.lampreydb.com), which includes detailed qualitative, quantitative, and spatial distribution information of each metabolite. Users can easily query and check their metabolites of interest, and/or identify unknown peaks from this database. In addition, a GitHub issue page (https://github.com/YonghuiDong/LampreyDB) is provided to allow users to report on errors, request new features, and/or submit new metabolite items. The current LampreyDB version includes information on over 1000 metabolites (2031 records from both positive and negative ion modes). We are now applying a fractionation-assisted NMR-based metabolomics approach to identify the unknown mass peaks from different lamprey tissues so that we can increase the number of metabolites and enhance the quality and reliability of the information in LampreyDB in the near future.

From a bloodsucker’s perspective, paradise is a place where the host blood does not clot, the blood flow is intense at the feeding site, and the host will not resist or harm the guests. While reality is different. The vertebrate hosts are equipped with three efficient weapons that fight against blood-sucking behaviors: hemostasis, inflammation, and immunity^53^. Blood-sucking animals have evolved many different strategies to succeed against all the complex barriers imposed by their hosts. Studies have shown that almost all bloodsucking arthropods have at least one anticlotting, one vasodilator, and one antiplatelet component, and in many cases, more than one substance is present in each category. In our study, we showed that in addition to relying on several active proteins and peptides, lampreys also secret many metabolites from their buccal glands to counteract the host responses and ensure successful and continuous blood-feeding. It has long been known that most KP metabolites are immune suppressors, and our study showed that kynurenine also functions as a vasodilator in lampreys. Although KP has been well-studied, one striking difference in our study is that the KP metabolites and the first enzyme TDO are found exclusively present in the buccal gland, which is different from reported in other animals and human that KP metabolites and TDO mostly exist in liver. This may imply a unique blood-sucking mechanism in lampreys. PGs are a group of well-known blood-sucking-related metabolites in other bloodsucking animals. They have been demonstrated to possess multiple roles to assist blood-feeding, such as increasing vasodilation, suppressing inflammation, and inhibiting wound healing. The identification of PGs in lampreys suggests that all bloodsuckers may share some similar blood-feeding strategies. These findings demonstrate the complex nature of lamprey buccal gland and highlight the diversity in the mechanisms utilized for blood-sucking in lampreys.

## Methods

### Chemicals

All chemicals and solvents were analytical or HPLC grade. Water, methanol, acetonitrile, formic acid were purchased from CNW Technologies GmbH (Düsseldorf, Germany). L-2- chlorophenylalanine was from Shanghai Hengchuang Bio-technology Co., Ltd. (Shanghai, China). L-tryptophan was purchased from Sangon (Shanghai, China), and L-kynurenine from Macklin (Shanghai, China).

### Lamprey model and ethical approval

The adult Japanese lampreys (*L. japonica*) at spawning migration stage were obtained in December 2020 in Songhua River in Heilongjiang province of China. Fourteen different lamprey tissues, i.e., heart, liver, kidney, brain, supraneural body, muscle, intestine, gill, eye, testis, ovary, buccal, blood, and notochord, were carefully dissected, and subjected to spatial metabolomics analysis.

The northeast lampreys (*Lampetra morii*) were obtained in December, 2020 in Yalu River in Liaoning province of China, and they were used for blood-sucking experiment. Sixty lampreys were randomly divided into six groups (10 in each group) and kept in fresh water at 10 ± 2 °C in dim light without feeding for 72 h. The buccal glands of three lamprey groups were dissected, and their secretion was collected before blood-sucking. Another three groups of lampreys were fed with catfish (*Silurus asotus*) for 20 min, and then their buccal gland secretion was collected. In addition, three sampling sites from the catfish, i.e., the blood-sucking site and two none-blood-sucking sites of the host, were collected. In total, the LCMS analyses include 5 sample groups: group 1 is the buccal gland from lamprey before blood-sucking; group 2 is the blood-sucking site from catfish; group 3 and 4 are none-blood- sucking sites from catfish, and group 5 is the buccal gland from lamprey after blood-sucking. The handling of lampreys was approved by the Animal Welfare and Research Ethics Committee of the Institute of Dalian Medical University (Permit number: AEE17013).

### Sample collection and LC-MS analysis

For metabolite extraction, 80 mg of the frozen fine power of each sample was extracted with 400 μL 80% methanol: water (v/v). 20 μL L-2-chlorophenylalanine (0.3 mg/mL) was added to each sample as internal standard. The extract was then briefly vortexed and sonicated at ambient temperature (25 ℃ to 28 ℃) for 10 min. Finally, the extracts were centrifuged at 13000 rpm, 4 ℃ for 15 min, filtered through a 0.22 μm PTFE filter (Acrodisc® CR 13mm; PALL), and transferred to LC vials for LCMS analysis. QC samples were prepared by mixing equal aliquots of all the samples.

A Dionex Ultimate 3000 RS UHPLC system fitted with Q-Exactive quadrupole- Orbitrap mass spectrometer equipped with heated electrospray ionization (ESI) source (Thermo Fisher Scientific, Waltham, MA, USA) was used for spatial metabolomics analysis in both positive and negative ion modes. An ACQUITY UPLC BEH C18 column (1.7 μm, 2.1 × 100 mm) was employed. The binary gradient elution system consisted of (A) water (containing 0.1 % formic acid, v/v) and (B) acetonitrile (containing 0.1 % formic acid, v/v) and separation was achieved using the following gradient: 5-20 % B over 0-2 min, 20-25 % B over 2-4 min, 25-60 B over 4-9 min, 60-100 % B over 9-14 min, the composition was held at 100 % B for 2 min, then 16-16.1 min, 100 % to 5 % B, and 16.1-18.1 min holding at 5 % B. The flow rate was 0.4 mL/min and column temperature was 45 ℃. All samples were kept at 4 ℃ during analysis. The injection volume was 5 μL. The mass range was from m/z 66.7 to 1000.5. The resolution was set at 70,000 for the full MS scans and 35,000 for HCD MS/MS scans. The Collision energy was set at 10, 20 and 40 eV. The mass spectrometer operated as follows: spray voltage, 3,000 V (+) and 2,500 V (−); sheath gas flow rate, 45 arbitrary units; auxiliary gas flow rate, 15 arbitrary units; capillary temperature, 350 °C. The QCs were injected at regular intervals (every 10 samples) throughout the analytical run to provide a set of data from which repeatability can be assessed.

### LCMS data processing

Raw data quality was first checked using R package RawHummus ^12^ on QC samples in both positive and negative ion modes. The resulting QC reports are shown in File S1-S2. Data pre-processing and metabolite identification were performed using three different software tools, i.e., Compound Discoverer (v.3.3, Thermo Scientific), Progenesis QI (v.2.3, Waters), and MS-DIAL (v.4.0) ^15, 16^. Of which, Progenesis QI and MS-DIAL were mainly used for metabolite identification purpose. For each data processing software, multiple parameters were tuned, and the optimized settings were summarized in Table S1-S3. The resulting feature tables (positive and negative ion mode) from Compound Discoverer were exported into csv format and subjected to subsequent statistical analysis using different R software packages. To compare the relative metabolite abundance among different tissue groups, one- way analysis of variance (ANOVA) was performed; If deemed significant (p-value < 0.05), post hoc multiple comparison analysis was performed with false discovery rate correction. Principal Component Analysis was performed with R package MSBox (https://cran.r-project.org/web/packages/MSbox/index.html); heatmap and hierarchical clustering analysis were performed using R package pheatmap (https://cran.r-project.org/web/packages/pheatmap/index.html). Other figures such as bar plots and line plots were produced using R package ggplot2 ^54^ and ggbreak ^55^. Data are represented as means ± SD of the mean (n = 3).

### Lamprey spatial metabolomics database construction

The LampreyDB database (https://www.lampreydb.com) was organized with MySQL (v.8.0) and Django (v.3.0.6). The web interface was developed using HTML with JavaScript. The interactive anatomy heatmap was produced with home written script and Python package beautiful soup (https://pypi.org/project/beautifulsoup4/). Other figures such as MS spectrum plots were produced using Python package plotly (https://plotly.com/python/). LampreyDB is hosted on Microsoft Azure cloud service.

### Gene cloning and bioinformatics analysis

The full-length open reading frames (ORF) of the *Lj-TDO* (tryptophan 2,3-dioxygenase), *Lj- AADAT* (aminoadipate aminotransferase), *Lj-IDO* (indoleamine 2,3-dioxygenase) and *Lj- KMO* (kynurenine monooxygenase) genes were obtained by PCR. Primer Premier 5.0 was used to design specific PCR primers in open reading frame. The sequences of all primers used for gene syntheses were listed in Table S4. The amino acid sequences of TDO, AADAT, IDO, and KMO used in sequence alignments and bioinformatics analysis were obtained from the NCBI database and saved in FASTA format. Sequence alignment was performed using Clustal W. The aligned sequences were used to construct phylogenetic trees using Neighbor-joining (NJ) method with 1000 bootstrapped replicates in the MEGA-X with the pairwise- deletion option. The conserved motifs were procured from online multiple expectation maximization (MEME, http://meme-suite.org/tools/meme) and the number of motifs was select: 15. The output search results were drawn with TBtools software. Conserved functional protein domains were predicted with Simple Modular Architecture Research Tool (SMART, http://smart.embl-heidelberg.de/). To better understand the evolution of gene family between jawless and jawed vertebrates, neighboring genes environment of TDO, AADAT, IDO, and KMO of representative animals were conducted by using the Genomicus online tool (https://www.genomicus.bio.ens.psl.eu/genomicus-92.01/cgi-bin/search.pl) and the Ensembl database (https://www.ensembl.org/index.html). The gene structure analysis was performed by using the Ensembl database. SWISS-MODEL (http://swissmodel.expasy.org/interactive) online was used to predict the 3D structure of Lj-TDO, Lj-AADAT, Lj-IDO and Lj-KMO.

### Real-time quantitative PCR analysis

The total RNA samples were isolated separately from the leukocytes, ovary, brain, livers, muscles, gills, oral glands, intestines, hearts, kidney, testis and supraneural bodies by using RNAiso reagent (TaKaRa, China). Leucocytes were collected from circulated blood. The cDNA was then synthesized using HiScript®II Q Select RT SuperMix for qPCR (+gDNA wiper) Kit (Vazyme, China) according to the manufacturer’s protocol. The specific primers for qPCR were designed using Primer Premier 5.0 program and the primer sequences were shown in Table S4. The qPCR was carried out in 20 µL reactions of 2×chamQ SYBR Color qPCR Master Mix (low ROX Rremixed) (Vazyme Biotech, China), forward primer (10 µM), reverse primer (10 µM) and cDNA 200 ng. Quantitative real-time PCR was performed with qTOWER 2.0 Real Time PCR System and the following program: 30 s at 95 ℃, 40 cycles of 10 s at 95 ℃ and 30 s at 60 ℃. The change in threshold cycle (ΔCT) was calculated by subtracting the average CT of *Lr-GAPDH* mRNA from the average CT of the target genes. All experiments were performed in triplicate.

### Vasomotor reactivity assessment

The vasomotor reactivity was measured following the protocol described elsewhere ^23^. In brief, catfish aortas were dissected, cut it into 1.8-2.0 mm aortic ring, and placed in a culture dish. The viability was then confirmed by incremental constriction to high potassium solutions (KCl) with the final concentration of 60 mmol/L. Aortic rings were next pre- incubated with 60 mmol/L KCl, pre-constricted to the maximum response, and then the blood vessel diameters were measured under a stereoscopic microscope. Finally, the vessel diameters were recorded 15 min after addition of different concentrations of kynurenine solutions (i.e., 2 mM, 4 mM, and 8 mM). PBS was added as the blank control group, and all the measurements were repeated five times (n = 5).

## Acknowledgments

This work was funded by Chinese National Natural Science Foundation Grant (No. 31772884, 32070518), Liaoning Climbing Scholar, the Distinguished Professor of Liaoning (No. XLYC2002093), the Project of the Educational Department of Liaoning Province (No. LJKZ0962), Open Fund of Key Laboratory of Biotechnology and Bioresources Utilization (Dalian Minzu University), and Ministry of Education (No. KF2022003).

## Contributions

M.Gou, Y. Pang and Y.Dong designed and directed the study. M.Gou, X.Duan, J.Li and Y.Wang performed all experiments. M.Gou and Y.Dong performed data analysis. M. Gou, X.Duan and Y.Dong prepared the LampreyDB database. Y.Dong wrote the draft. All other authors reviewed and edited the manuscript.

## Data Availability

The sequences files used in this study are available in the National Center for Biotechnology Information (accession number: ON814546, ON814547, ON814548 and ON814549). The metabolomics data have been archived in MetaboLights with MTBLS5857.

## Table of Content

1. Table S1. The optimal set of parameters for Compound Discoverer
2. Table S2. The optimal set of parameters for Progenesis QI
3. Table S3. The optimal set of parameters for MS-DIAL
4. Table S4. The primer sequences used in the study
5. Fig. S1 Head-to-tail plot of experimental and library ESI-MS/MS spectra of tryptophan.
6. Fig. S2 Head-to-tail plot of experimental and library ESI-MS/MS spectra of L-kynurenine.
7. Fig. S3 Head-to-tail plot of experimental and library ESI-MS/MS spectra of L-kynurenic acid.
8. Fig. S4 Head-to-tail plot of experimental and library ESI-MS/MS spectra of 3- hydroxyanthranilic acid.
9. Fig. S5 Head-to-tail plot of experimental and library ESI-MS/MS spectra of xanthurenic acid.
10. Fig. S6 Head-to-tail plot of experimental and library ESI-MS/MS spectra of anthranillic acid.
11. Fig. S7 Head-to-tail plot of experimental and library ESI-MS/MS spectra of prostaglandin E2.
12. Fig. S8 Head-to-tail plot of experimental and library ESI-MS/MS spectra of prostaglandin E3.
13. Fig. S9 Head-to-tail plot of experimental and library ESI-MS/MS spectra of prostaglandin F2α.
14. Fig. S10 Head-to-tail plot of experimental and library ESI-MS/MS spectra of prostaglandin J2.

**Table S1.**
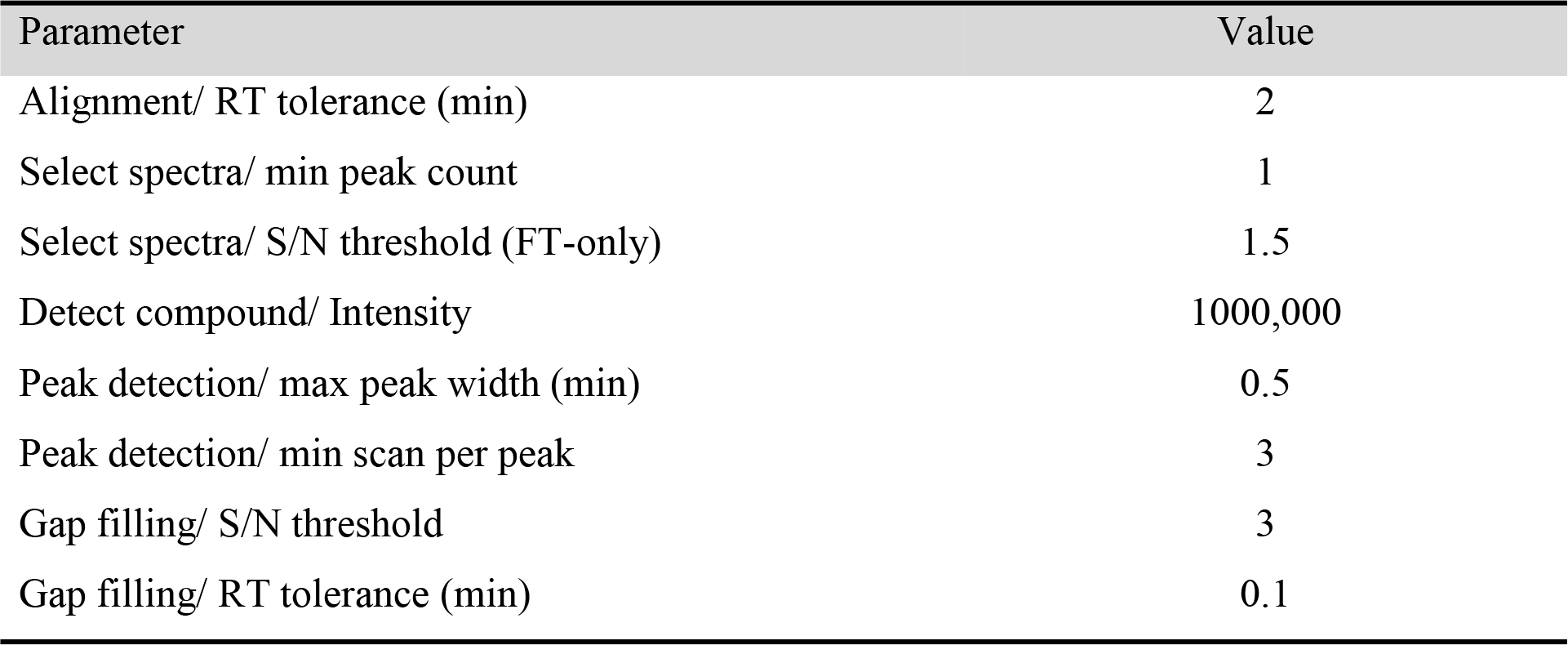
The optimal set of parameters for Compound Discoverer

**Table S2.**
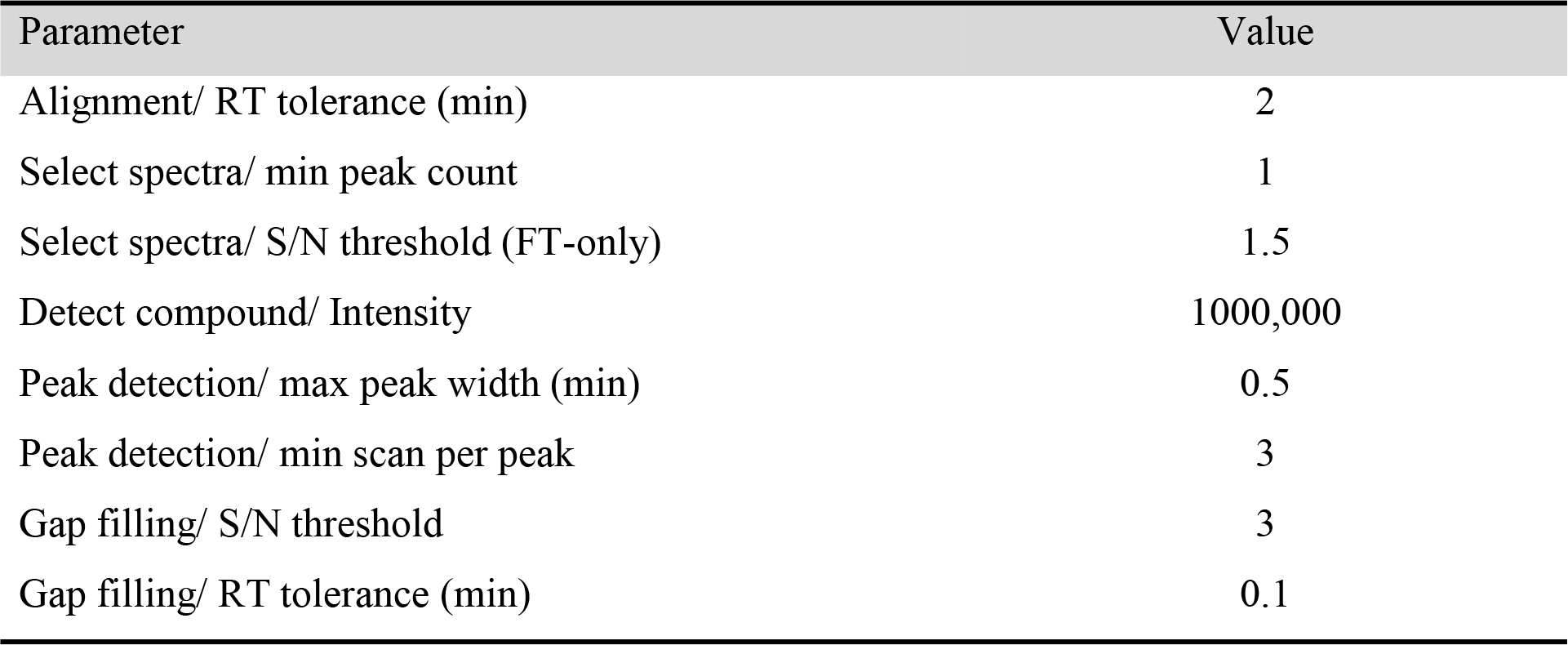
The optimal set of parameters for Progenesis QI

**Table S3.**
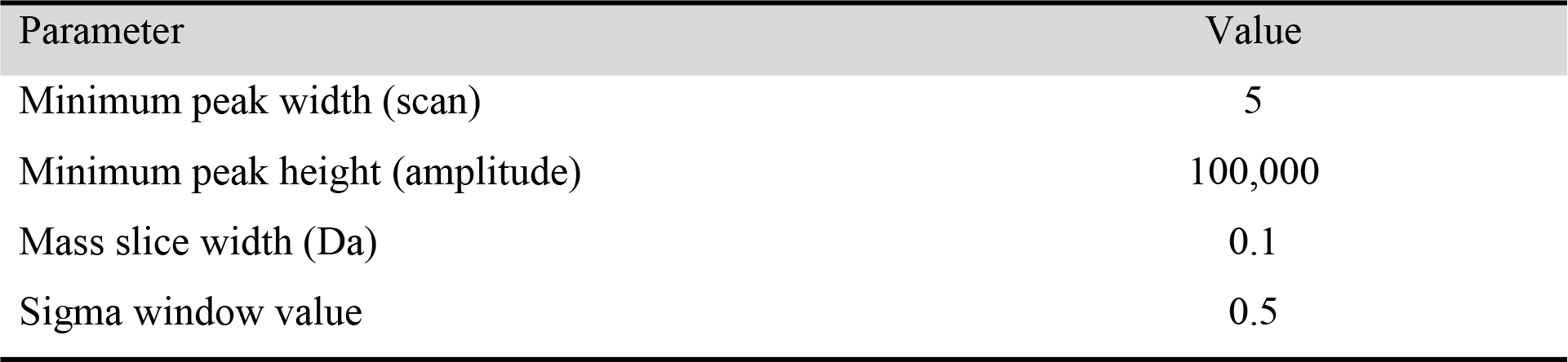
The optimal set of parameters for MS-DIAL

**Table S4.**
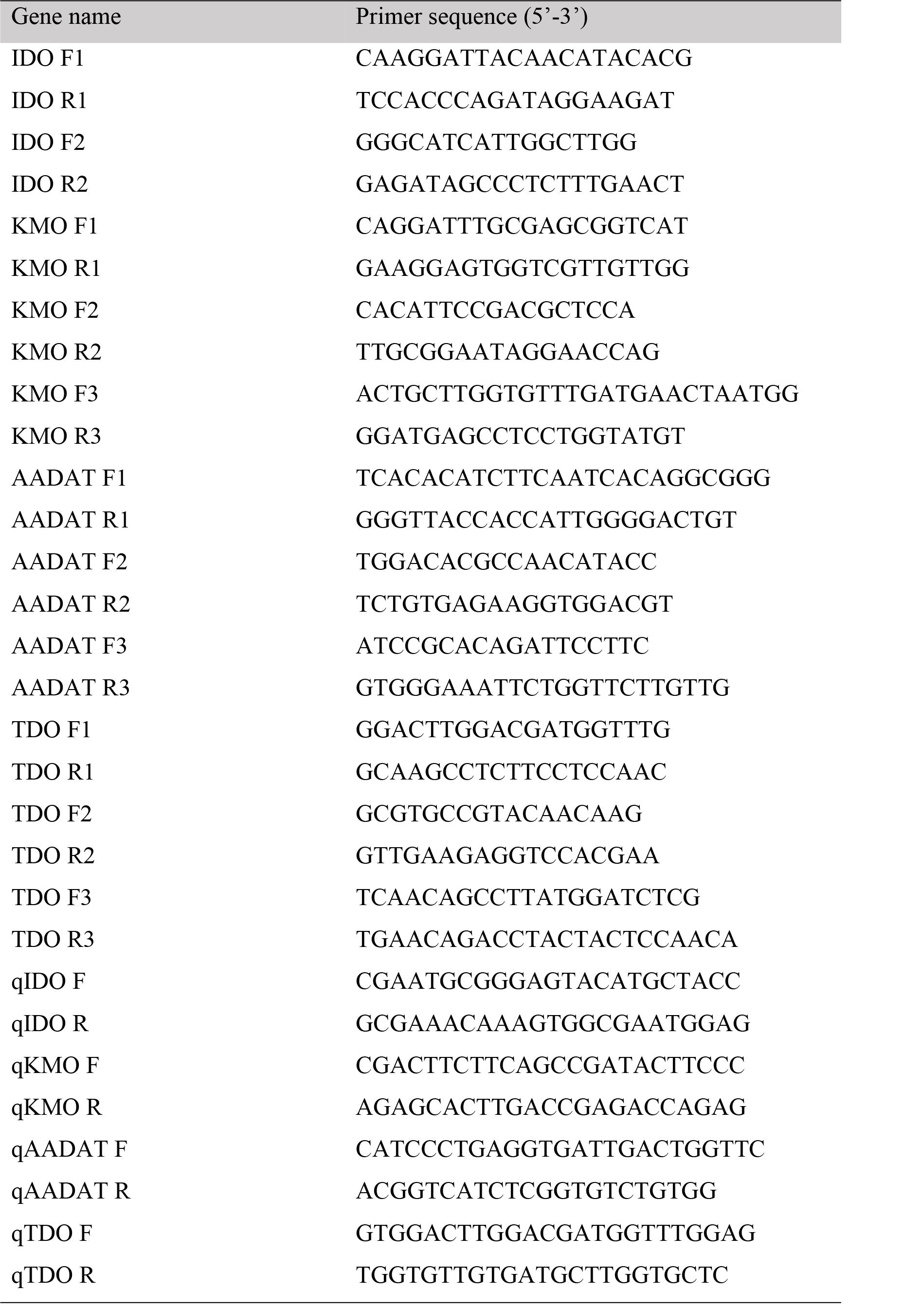
The primer sequences used in the study

**Fig. S1.**
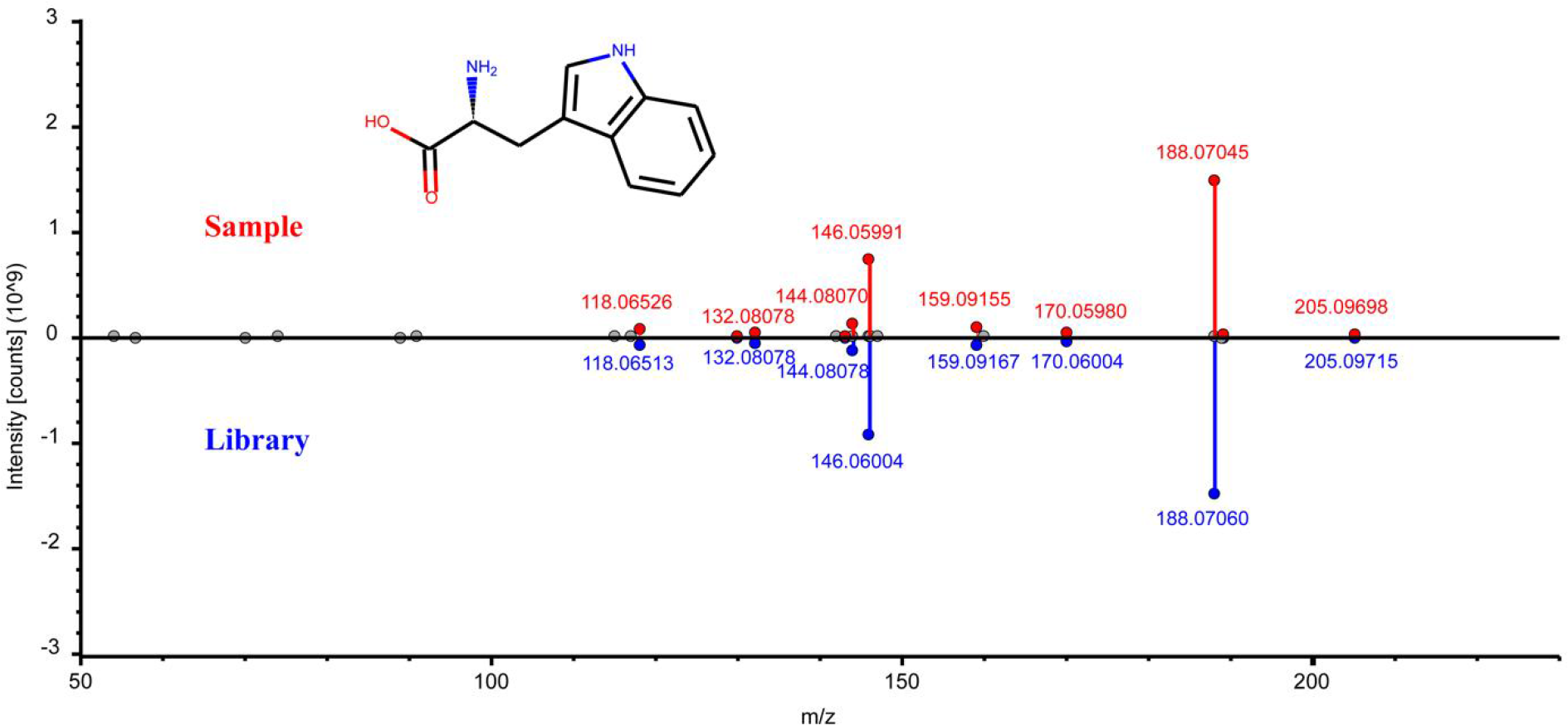
Head-to-tail plot of experimental and library ESI-MS/MS spectra of tryptophan.

**Fig. S2.**
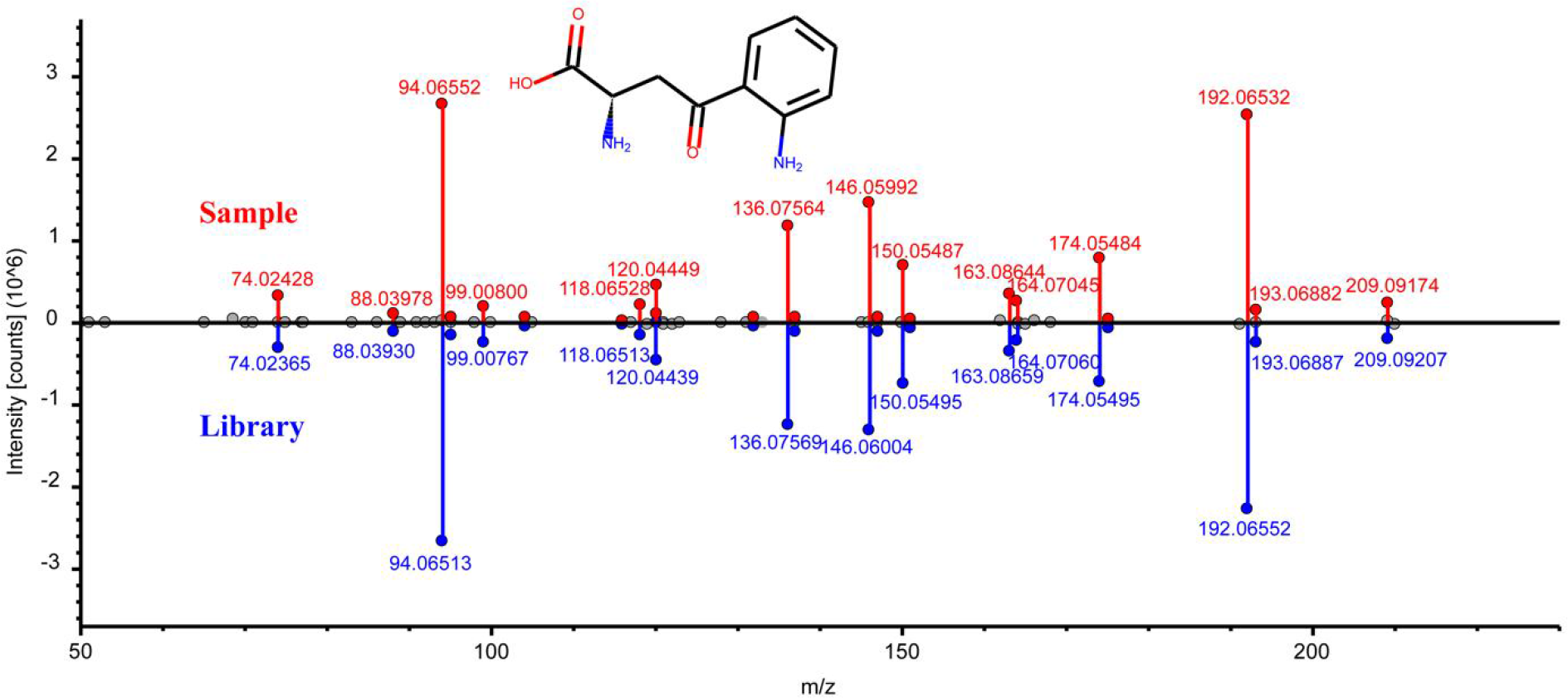
Head-to-tail plot of experimental and library ESI-MS/MS spectra of L-kynurenine.

**Fig. S3.**
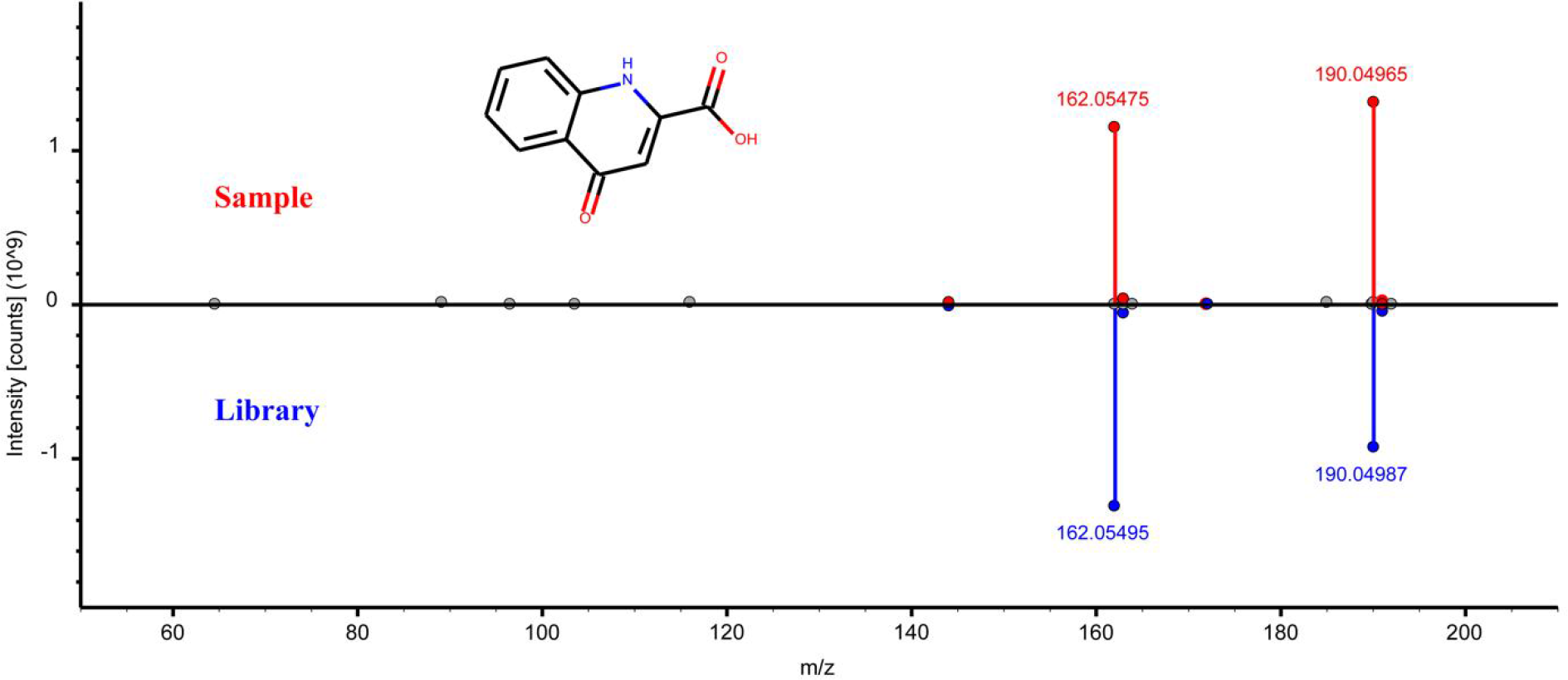
Head-to-tail plot of experimental and library ESI-MS/MS spectra of L-kynurenic acid.

**Fig. S4.**
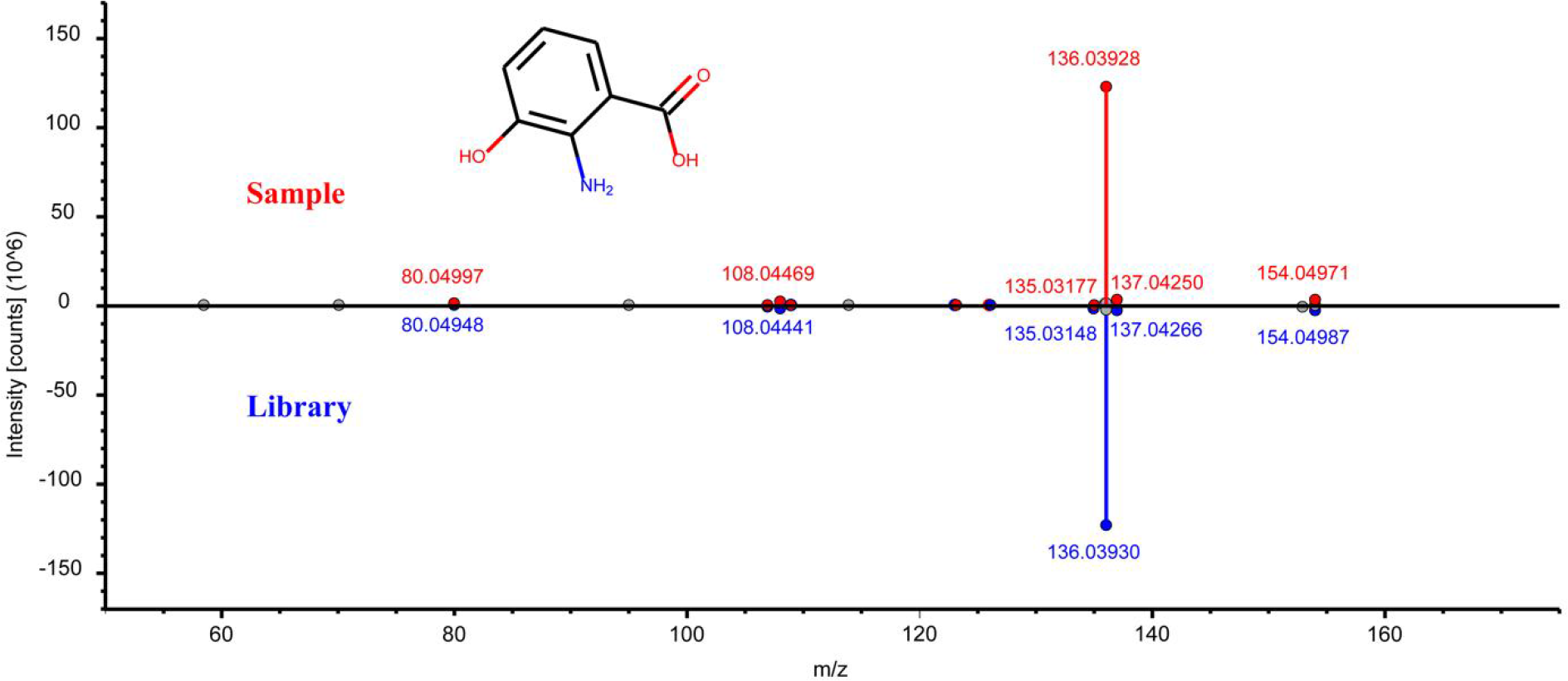
Head-to-tail plot of experimental and library ESI-MS/MS spectra of 3- hydroxyanthranilic acid.

**Fig. S5.**
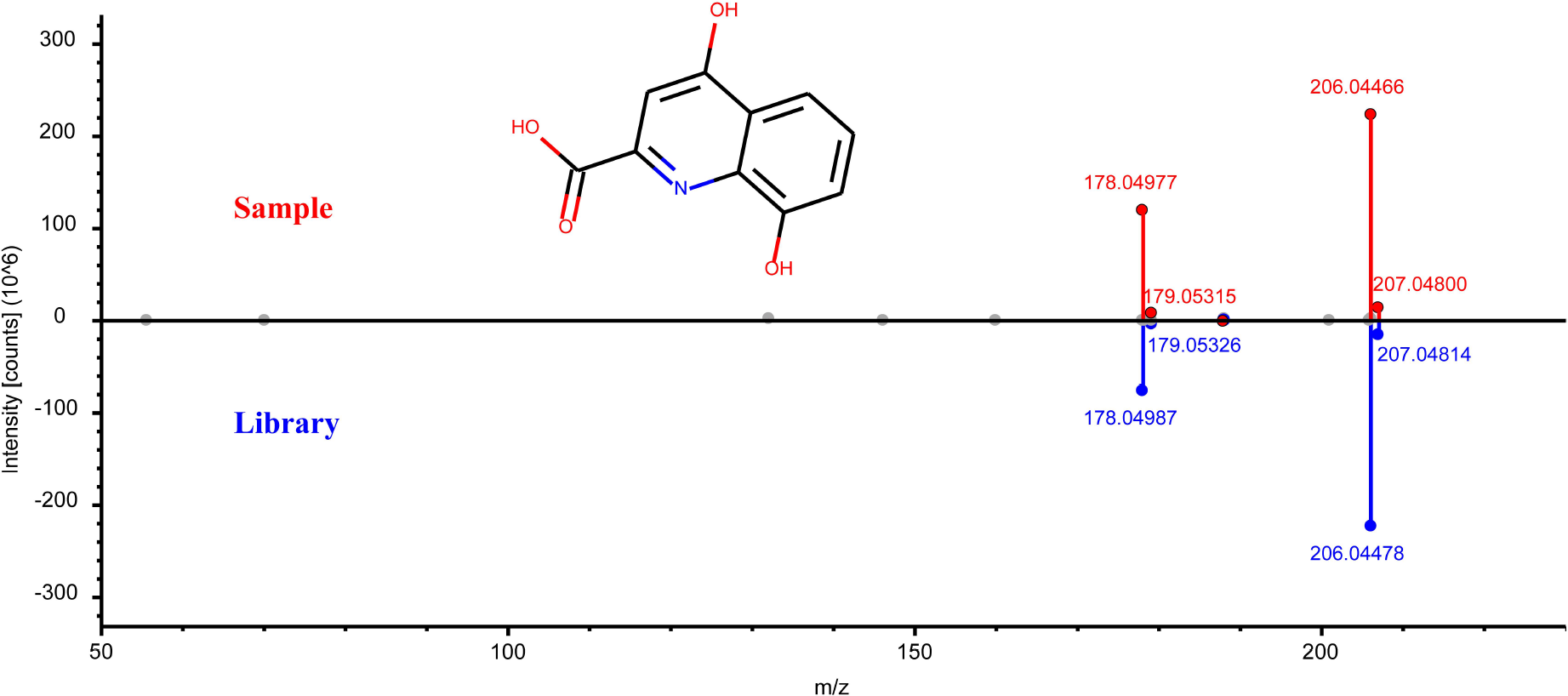
Head-to-tail plot of experimental and library ESI-MS/MS spectra of xanthurenic acid.

**Fig. S6.**
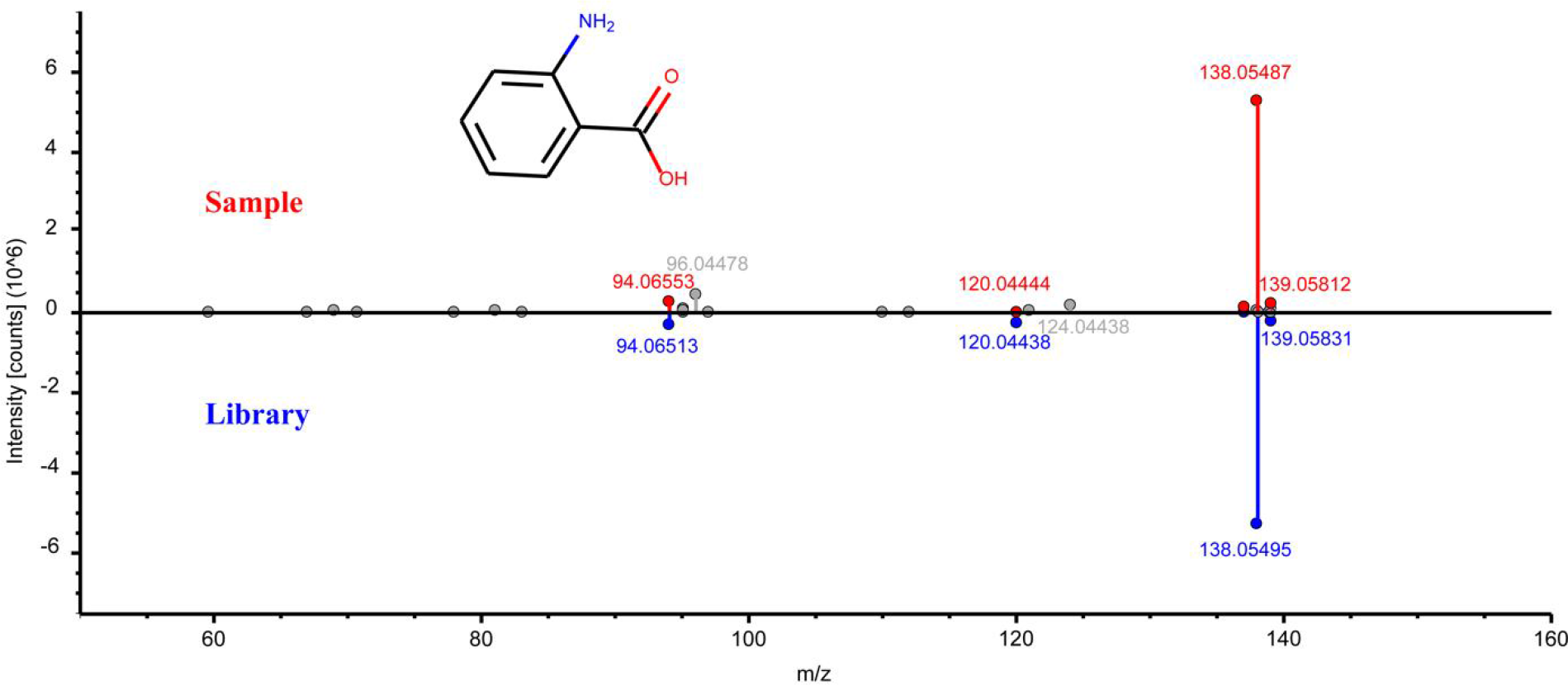
Head-to-tail plot of experimental and library ESI-MS/MS spectra of anthranillic acid.

**Fig. S7.**
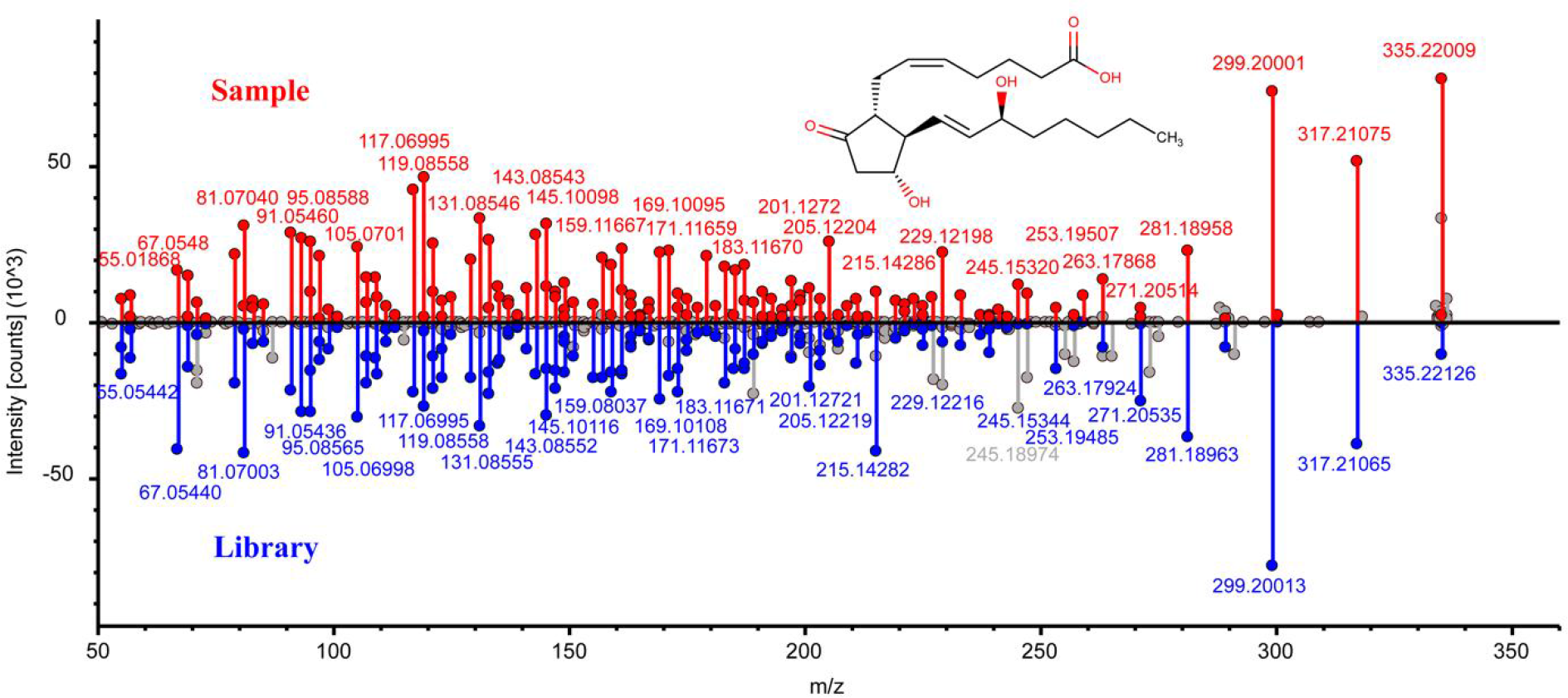
Head-to-tail plot of experimental and library ESI-MS/MS spectra of prostaglandin E2.

**Fig. S8.**
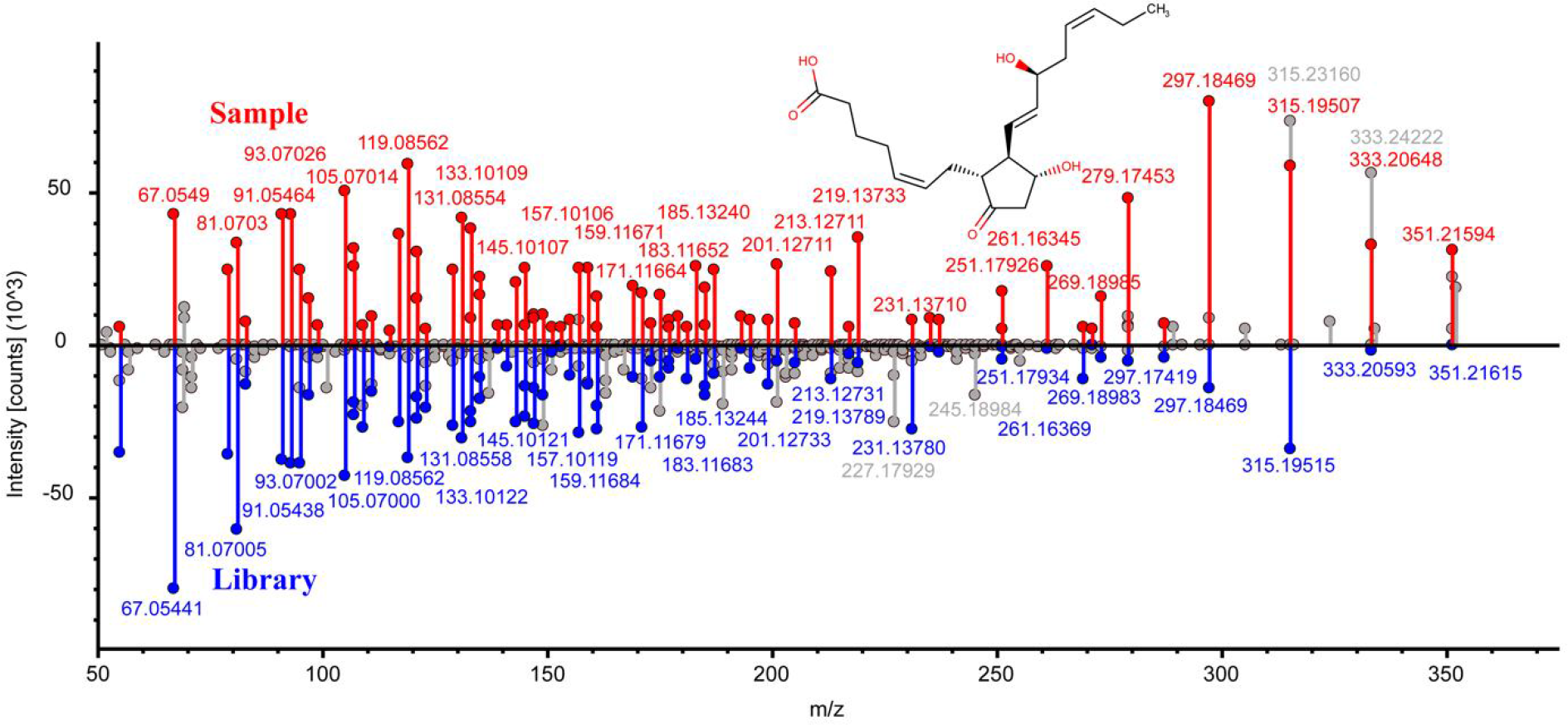
Head-to-tail plot of experimental and library ESI-MS/MS spectra of prostaglandin E3.

**Fig. S9.**
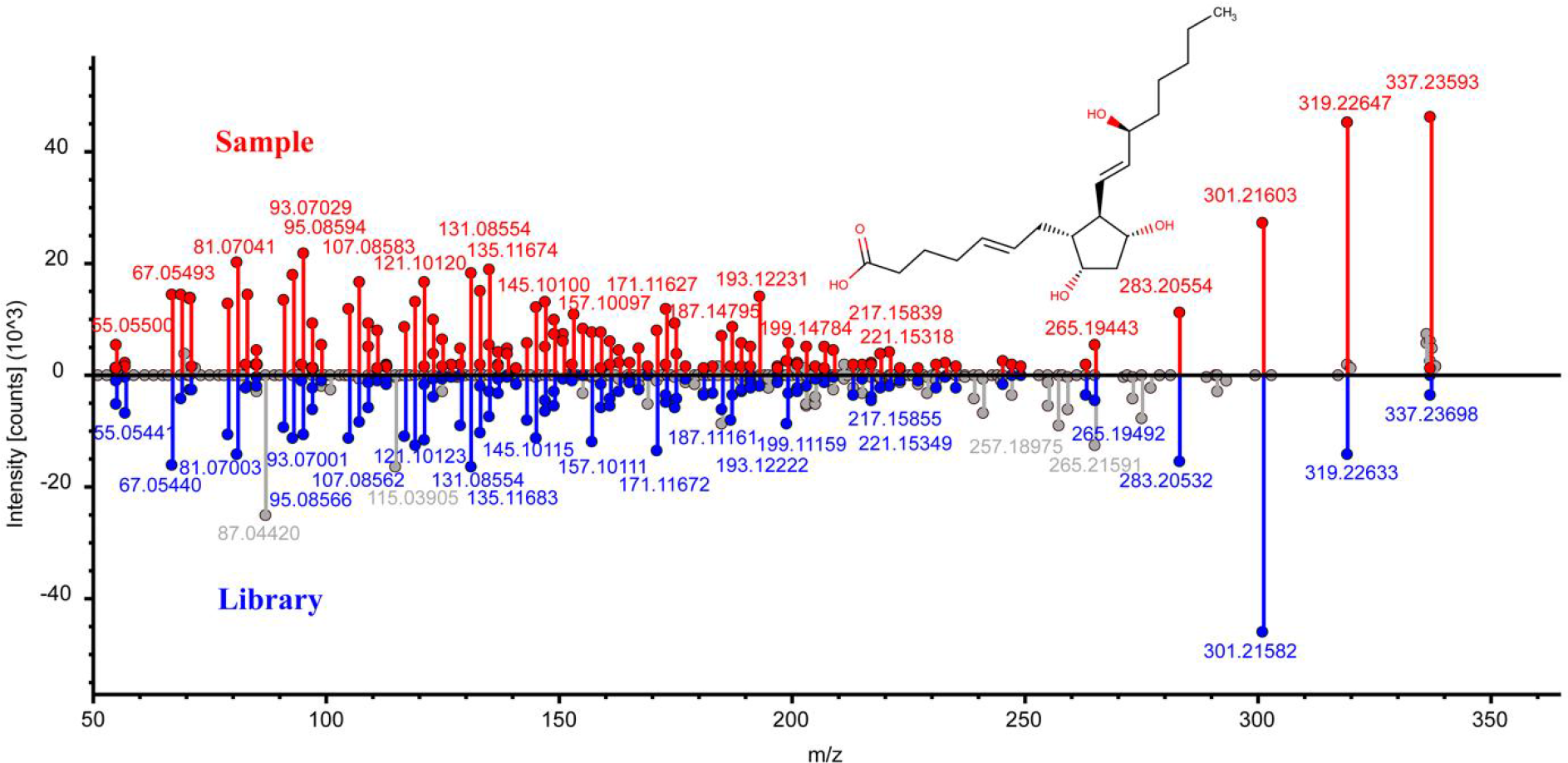
Head-to-tail plot of experimental and library ESI-MS/MS spectra of prostaglandin F2α.

**Fig. S10.**
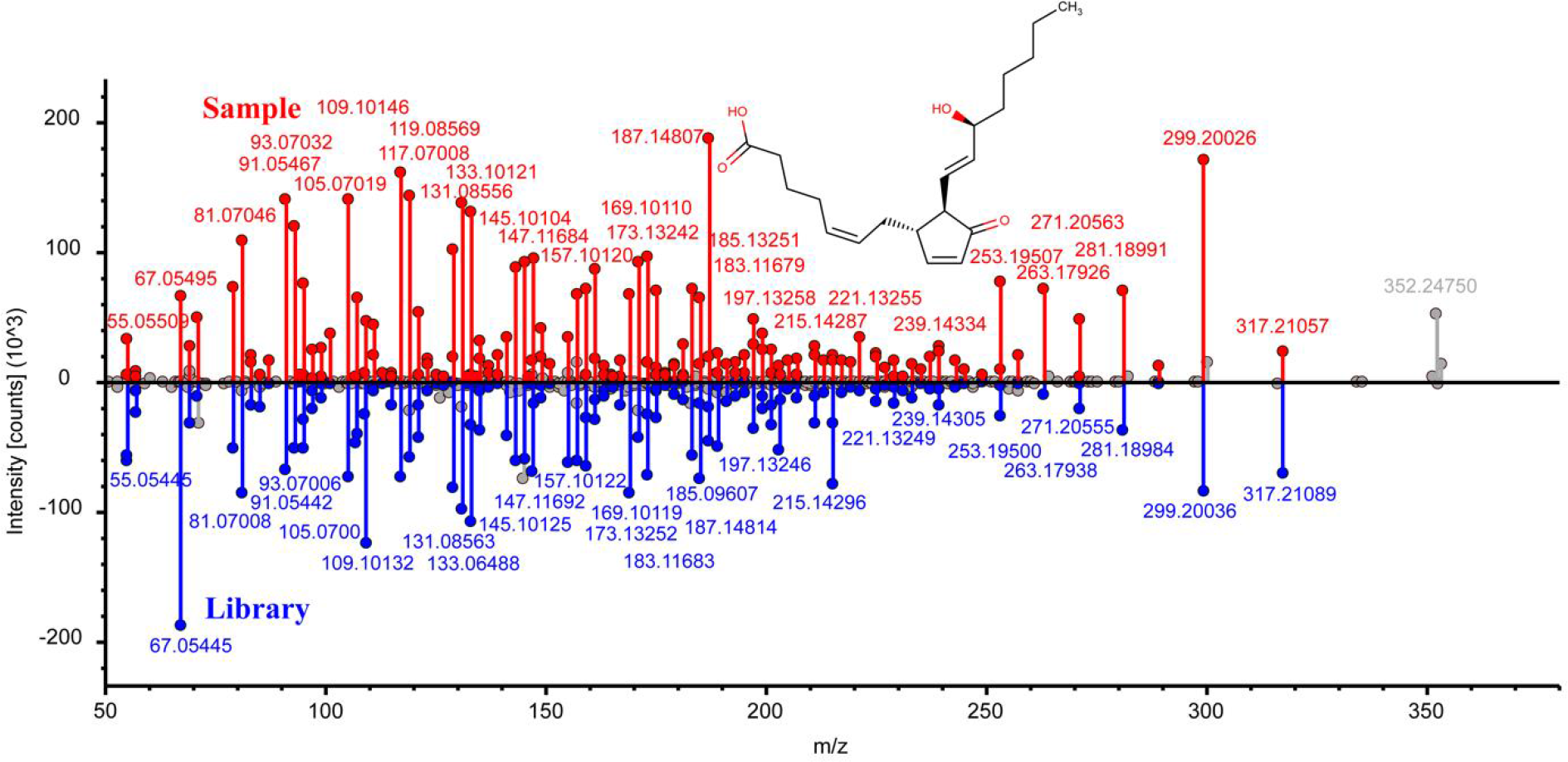
Head-to-tail plot of experimental and library ESI-MS/MS spectra of prostaglandin J2.

**Fig. S11.**
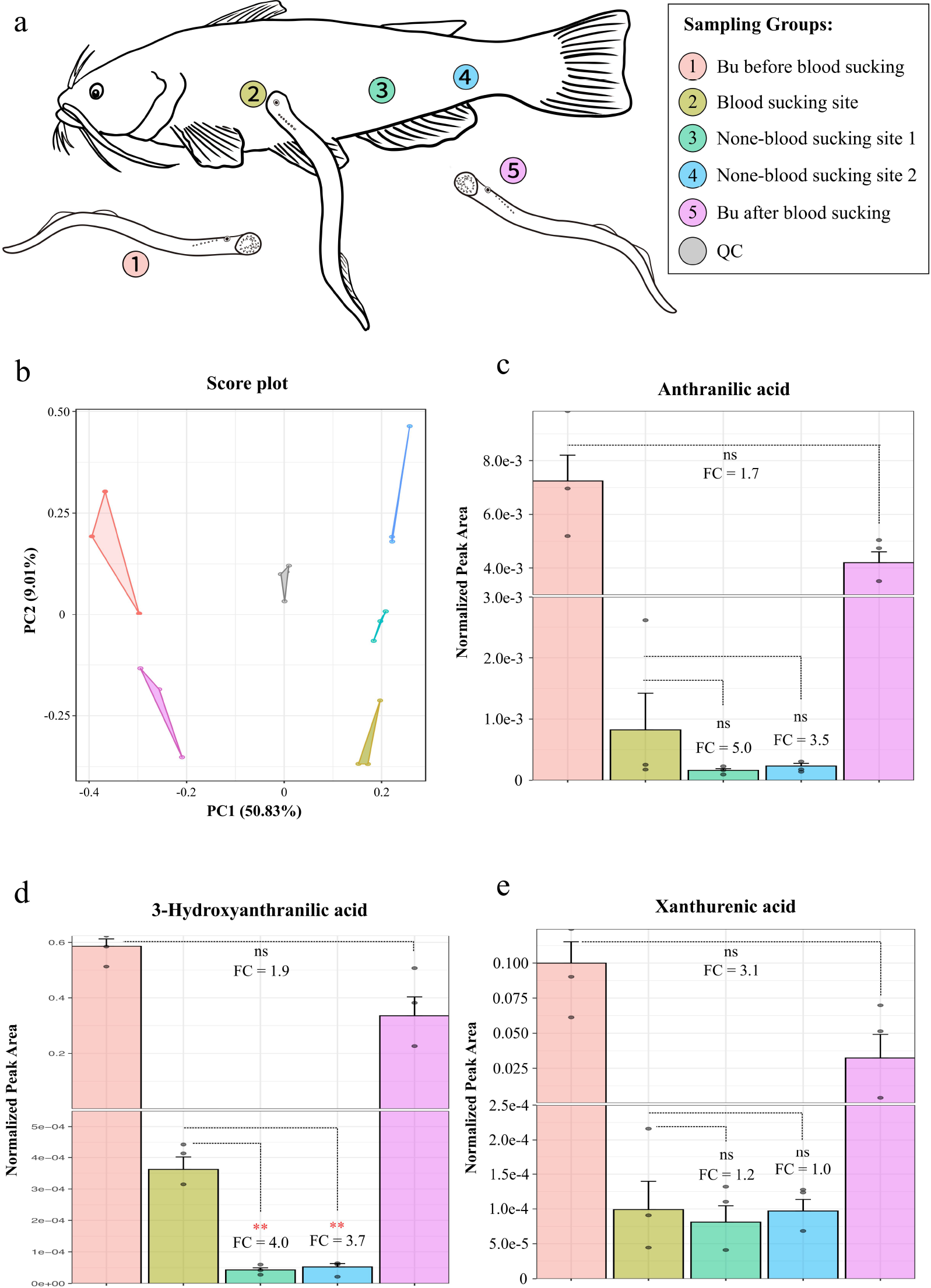
Experimental design for lamprey blooding sucking. **a**, Five groups of lampreys were used for blood-sucking experiment. Group 1 is the buccal gland collected from lampreys before blood-sucking; Group 2 is the feeding-site from the host fish (catfish); Groups 3 and 4 are non-feeding sites from the host fish; and group 5 is the buccal glands collected from lampreys after blood-sucking. **b**, Principal component analysis (PCA) score plots of the metabolic profiles of the 5 sample groups (n = 3 for each group). **c-e**, Measurement of relative peak areas of anthranilic acid (c), 3-hydroxyanthranilic acid (d), xanthurenic acid (e). The data are shown as the mean ± SD (n = 3). Asterisks denote significant differences (ns, not significant; *, p < 0.05; **, p < 0.01).

**Fig. S12.**
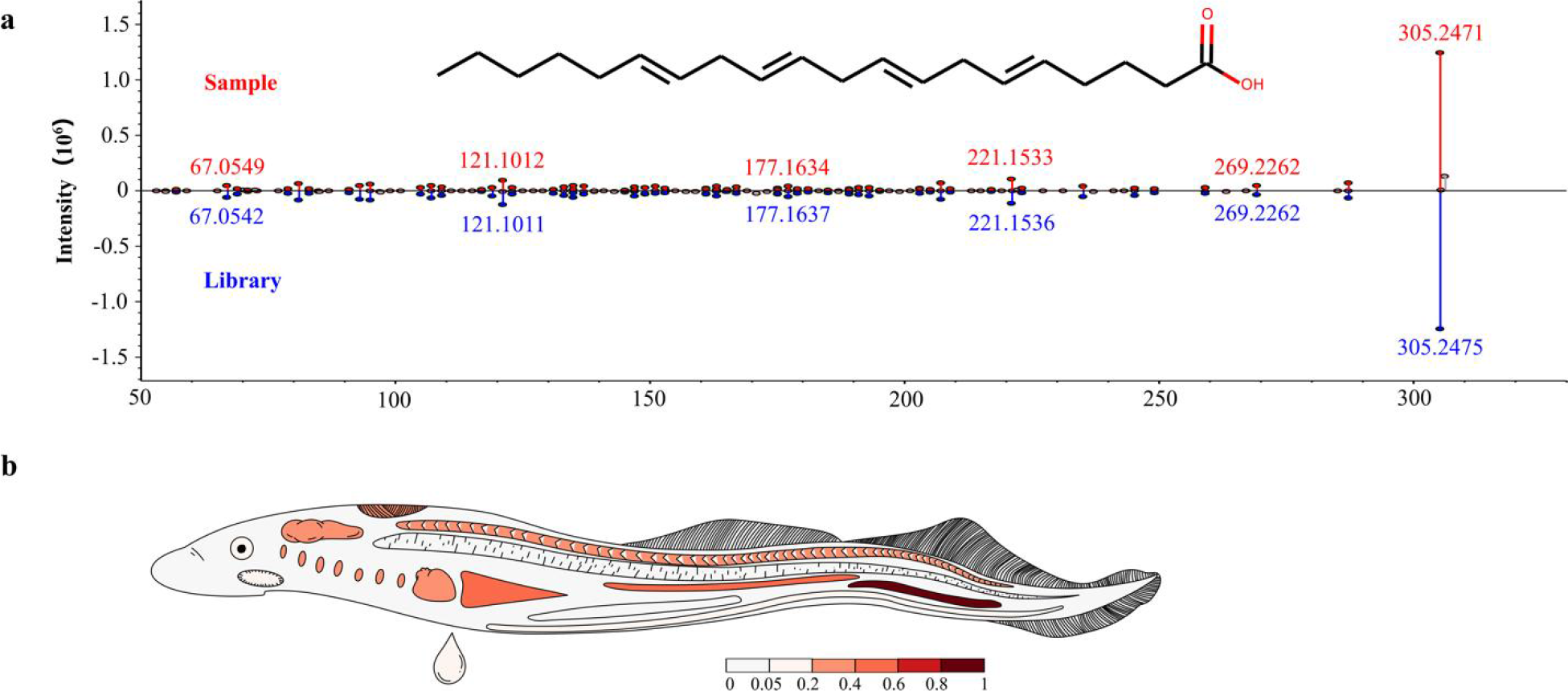
Identification of arachidonic acid in lampreys. **a,** Head-to-tail plot of experimental and library ESI-MS/MS spectra of arachidonic acid. **b**, Spatial distribution of arachidonic acid in lampreys.

**Fig. S13.**
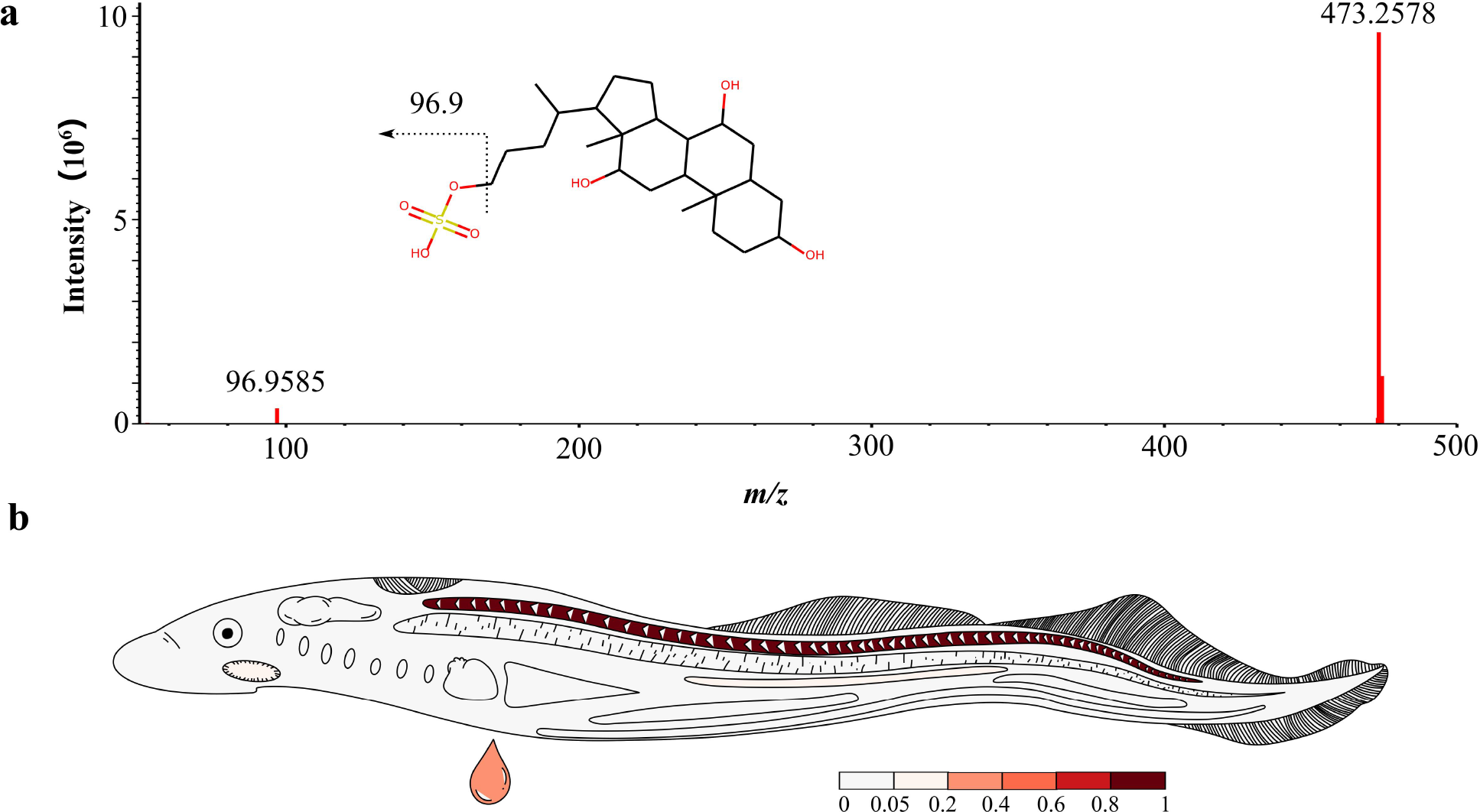
Identification of petromyzonol sulfate in lampreys. **a,** Head-to-tail plot of experimental and library ESI-MS/MS spectra of petromyzonol sulfate. **b**, Spatial distribution of petromyzonol sulfate in lampreys.

